# Enzyme Bioink for the 3D Printing of Biocatalytic Materials

**DOI:** 10.1101/2024.02.04.577699

**Authors:** Luca A. Altevogt, Rakib H. Sheikh, Thomas G. Molley, Joel Yong, Kang Liang, Patrick Spicer, Kristopher A. Kilian, Peter R. Wich

## Abstract

The field of 3D biofabrication faces major challenges on the road to printing fully functional tissues and organs. One of them is adding functionality to the newly formed tissue for replicating an active biochemical environment. Native extracellular matrices sequester numerous bioactive species, making the microenvironment biochemically active. On the other hand, most 3D-printed constructs have limited activity, serving merely as mechanical scaffolding. Here we demonstrate active scaffolding through the integration of biocatalytic enzymes within the bioink. Enzymes are an attractive class of biocompatible and substrate-specific bioactive agents that can improve tissue regeneration outcomes. However, the difficulty in the application remains in providing enzymes at the targeted site in adequate amounts over an extended time.

In this work, a durable biocatalytic active enzyme bioink for 3D extrusion-based bioprinting is developed by covalently attaching the globular enzyme horseradish peroxidase (HRP) to a gelatin methacrylate (Gel-MA) biopolymer scaffold. Upon introducing methacrylate groups on the surface of the enzyme, it undergoes photo-crosslinking in a post-printing step with the methacrylate groups of Gel-MA without compromising its activity. As a result, HRP becomes a fixed part of the hydrogel network and achieves higher stability inside the gel which results in a higher concentration and catalytic activity for a longer time than solely entrapping the protein inside the hydrogel. We also demonstrate the cytocompatibility of this enzyme bioink and show its printing capabilities for precise applications in the field of tissue engineering. Our approach offers a promising solution to enhance the bioactive properties of 3D-printed constructs, representing a critical step towards achieving functional biofabricated tissues.

## 1. Introduction

Three-dimensional (3D) bioprinting has rapidly evolved in the field of tissue engineering in the past few years. The technology enables the precise positioning of cell-laden material for the fabrication of complex structures with well-defined three-dimensional architecture. The aim to create and replace fully functional tissues^1^ still faces many challenges such as scalability, preventing immune responses, maintaining high cell viability and function, vascularization, and creating biocompatible bioinks with both good mechanical properties and printability.^2, 3^ The key to solving this requires advanced fabrication methods and multifunctional bioinks as the main supporting material in the printing process.^4, 5^ These materials are commonly physically or chemically crosslinked hydrophilic polymer networks. Due to their ability to absorb high amounts of water and resemble native biological tissue, these hydrogels have been an excellent choice as materials for 3D bioprinting and scaffold biofabrication.^5, 6^ They provide physical integrity such as mechanics and regulation of porosity which enables nutrient and cell diffusion.^7, 8^ One of the most frequently used hydrogel materials for bioink formulations is gelatin (Gel), a denatured form of collagen.^9, 10^ It shows high biocompatibility, water solubility, facile preparation, relatively low cost, and thermo-reversible sol-gel transition to form physically crosslinked hydrogels at low temperatures.^11^ However, to achieve long-term mechanical stability, functionalization with methacrylate groups on the amino residues of the polymer backbone is usually done (Gel-MA). This enables covalent bonds between adjacent polymer chains in a photo-crosslinking step to create a durable network of the printed structure.^12^ The ability for physical or chemical functionalization can additionally lead to better adhesion^13^, can control sol-gel transition, create shape memory or develop self-healing behaviors in response to an external stimulus.^14^

Equally important to mechanical integrity to support tissue formation is the introduction and preservation of biological activity and tissue-specific functions within the hydrogel.^2, 15^ Bioactive compounds, like growth factors, are needed to promote various cellular functions, such as cell proliferation, differentiation, or ECM production and induce tissue integration.^2, 16^ The vascular endothelial growth factor (VEGF) is a common bioactive molecule that is incorporated into polymer scaffolds to provide a controlled release of angiogenic signals to encourage vascularization.^17^ Similarly, other compounds are added to go beyond the preservation of basic tissue functions, including facilitating wound healing processes, preventing microbial infection, or acting as diagnostic systems.^18–20^ For example, antibacterial agents like silver nanoparticles and antibiotics are incorporated to combat bacterial infection,^21^ or antibodies and enzymes can be used as biosensors, exemplified by the use of glucose oxidase as glucose detector.^22–24^

One type of bioactive molecules that has received less attention as a building block are enzymes. They represent a promising class of bioactive materials for applications in biocatalytic reactors,^25–27^ sensors^23, 28–30^ but also as an additive to aid tissue engineering.^18^ They are ideal candidates for adding functionality to biocatalytic hydrogels, due to their biocompatibility and high substrate specificity and selectivity, respectively. Examples of enzyme-based functionalities include the support of wound healing processes by quenching reactive oxygen species (ROS) and oxygen generation,^31–33^ the prevention of microbial infections,^21, 34^ and improved wound repair by controlled degradation of wound dressings.^18, 35^ Notably, in all mentioned studies, enzymes were physically entrapped or co-incubated within hydrogels. This can result in uneven enzyme distribution and low retention within biomaterial scaffolds due to enzymes leaching out, leading to short-term effects only.^31, 36^ Hence, attempts have recently been made to covalently attach enzymes inside hydrogels,^37, 38^ e.g. the conjugation of methacrylated lysozyme to chitosan^35^ and hydrogels of photocurable bovine serum albumin (BSA).^39^ However, these methods are restricted to mold-based casting methods and are not applicable to 3D bioprinting.

In the field of tissue biofabrication, enzymes have predominantly been used as crosslinkers to catalyze the formation of covalent bonds between polymers.^40–42^ Only a few examples describe the physical entrapment of enzymes within 3D-printed hydrogels.^43–47^ This includes the encapsulation of glucose oxidase and horseradish peroxidase within photocurable polyethylene glycol hydrogels as chemiluminescent biosensors of glucose,^23^ and the entrapment of alkaline phosphatase and thrombin within a poly(ethylene glycol) diacrylate (PEGDA) bioink for improved tissue formation.^48^ Similarly, BSA was immobilized in a stereolithography approach where its methacrylated form was crosslinked with PEGDA. However, during an additional thermal post-print curing procedure, the proteins denatured.^49^

While these examples show that proteins and enzymes have been incorporated in hydrogels and 3D printing materials, the challenge of efficient protein retention remains. It can be improved by increasing the density of gel networks, however, this also limits cell migration and proliferation.^50^ To circumvent this, a mild covalent attachment method that maintains enzymatic activity while also being suitable for 3D bioprinting is required.

In this work, we developed the first functional enzyme bioink by covalently linking the antioxidant enzyme horseradish peroxidase (HRP) to a gelatin biopolymer scaffold. Both the biopolymer and the enzyme were modified with methacrylate groups to form stable covalent connections during post-print photocuring. The mild and biocompatible crosslinking method preserves the functional and structural activity of the enzyme, while at the same time increasing its retention within the gel. The resulting bioink can be used in standard extrusion-based 3D printing methods and provides the opportunity to obtain functional hydrogels with intrinsic biocatalytic activity. Overall, this work highlights a versatile approach of conjugating enzymes to bioinks, providing a new methodology to create functional and bioactive matrices for biofabrication and tissue engineering.

## 2. Results

### 2.1 Material Synthesis

All bioink components were prepared separately, starting with the underlying polymer scaffold. For this, gelatin was chosen due to its facile preparation and common application in the field of 3D extrusion-based bioprinting.^51^ To enable photo-crosslinking for post-print stability, free amine groups on gelatin were functionalized with methacrylate groups using a one-pot synthesis.^52, 53^ ^1^H-NMR and fluorescamine assays demonstrate a degree of substitution (DS) of 98%, providing a high amount of possible crosslinking locations (Figures S1 and S2).

To conjugate HRP to Gel-MA, methacrylate functional groups were introduced at free amino groups of lysines on the surface of the enzyme, similar to modifications previously performed on bovine serum albumin (BSA) for the generation of photocurable hydrogels.^39, 49^ BSA represents a robust and relatively large globular protein with 30-35 accessible amino groups. In contrast, enzymes, feature well-designed active sites, either on the surface or buried as deep binding pockets. Hence, substrate accessibility and structural integrity after the chemical modification play a more important role. HRP is a relatively small protein with a molecular weight of 44.1 kDa and represents a glycoprotein that carries heme as a redox cofactor in the core area. Previous work has shown that HRP is relatively robust^54^ and can be conjugated with biopolymers.^54^ Here, we chose HRP as proof-of-principle enzyme due to the easy biotechnological detection of its catalytic activity with colorimetric assays,^55, 56^ and its ability to scavenge and detect harmful reactive oxygen species (ROS) like H_2_O_2_.^57, 58^ In addition, its relatively small physical size is ideal for probing the efficiency of a covalent attachment to a bioink scaffold, enabling easy distinction between sole physical entrapment and covalent anchoring. Since HRP only presents 6 free amino residues on its surface,^59^ we aimed to introduce as many methacrylate groups as possible.

For the modification with methacrylic anhydrate (MAA), a carbonate-bicarbonate at pH 9 was chosen as enzyme-compatible buffer to achieve a high buffer capacity and increase the reactivity of the surface amines. Additionally, reaction time, temperature, and the ratio of MAA/lysine were optimized to achieve a high degree of substitution of methacrylate groups (Figure 1B).

**Figure 1.**
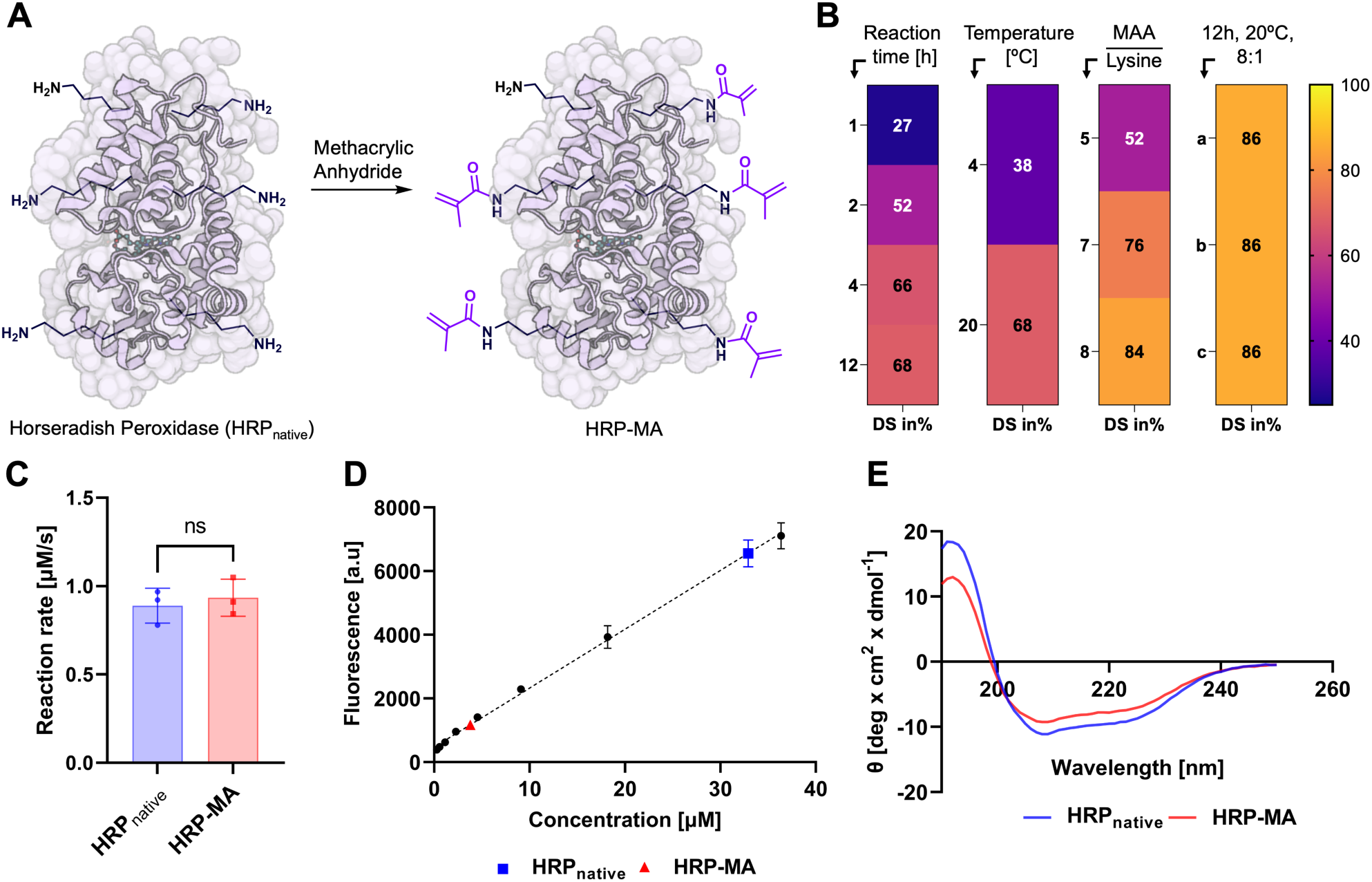
(A) Functionalization of the enzyme HRP with methacrylate groups. (B) Optimization of the methacrylation process in terms of reaction time (20 °C, 5:1 MAA/lysine), reaction temperature (12 h, 5:1 MAA/lysine), MAA/lysine ratio (12 h, 20 °C) and reproducibility (12 h, 20 °C, 8:1 MAA/Lysine). (C) ABTS enzyme activity assay of HRP_native_ vs HRP-MA; one-way ANOVA ns P > 0.05. (D) Fluorescamine assay to determine free amino groups on the surface of HRP_native_ vs HRP-MA. (E) Circular dichroism (CD) spectrum to determine changes in the secondary structure of HRP-MA compared to HRP_native_.

**Figure 2.**
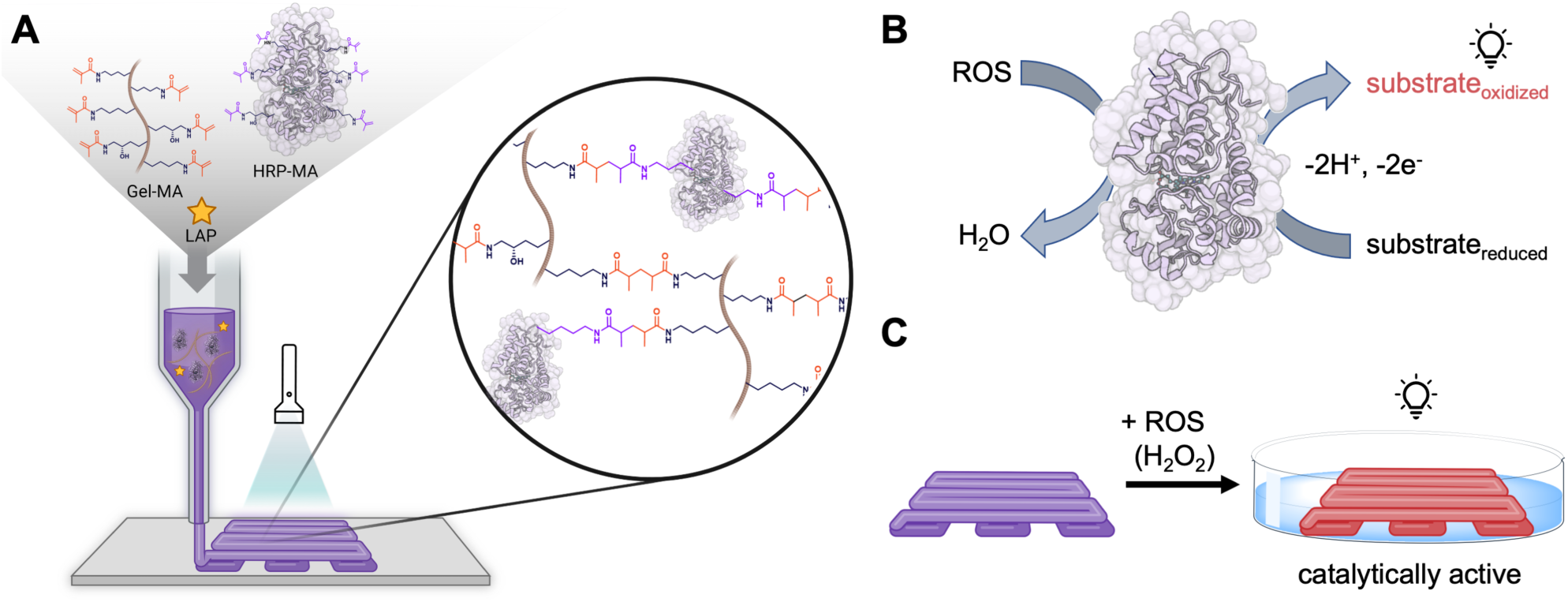
Overview of enzyme bioink preparation and catalytic activity. (A) 3D extrusion-bioprinting of enzyme bioink components and photo-crosslinking of enzymes with biopolymers. (B) HRP can catalytically transform reactive oxygen species (ROS) into water while oxidizing a colorimetric substrate. (C) The covalent incorporation of enzymes inside the bioink allows the formation of printed structures that are biocatalytically active and e.g. can scavenge harmful ROS.

The optimized HRP-MA functionalization protocol starts with the dissolution of HRP in CB buffer followed by the addition of a 0.1 w/v% of a stock solution of MAA in DMSO for a targeted MAA/lysine ratio of 8:1. The mixture was then stirred at room temperature for 12 hours, purified via centrifugation filtration, and lyophilized overnight. Successful methacrylation was further verified via fluorescamine assays and FTIR (Figures 1D, S2, S3). For this, free amino groups on HRP-MA were determined and compared with the number of free amino groups of non-functionalized HRP_native_. The analysis showed that HRP-MA has around 86% fewer free amines on their surface which means an average of 5 out of 6 amino groups were functionalized with a methacrylate group.

Circular dichroism (CD) spectroscopy (Figure 1E) showed only minimal changes in the secondary structure of HRP-MA after the modification process. Additional ABTS activity assays (Figure 1C) confirmed no significant difference in the activity of the modified enzyme compared to native HRP.

### 2.2 Enzyme Bioink Preparation

For the assembly of the final enzyme bioink, Gel-MA and HRP-MA were mixed with a photoinitiator in preparation for the hydrogel printing. Lithium acylphosphinate (LAP) was chosen as photoinitiator due to its absorption in the low-energy ultraviolet (UV) light range (370 nm) and low toxicity.^60^ For all samples, a stock solution of 11 w/v % Gel-MA was prepared at 40 °C. Next, LAP (2.5 mg/mL) and enzyme (0.625 mg/mL) were added in absence of light. To avoid contamination, the hydrogel was filtered once with a syringe filter (33 mm, Ø 0.45 *µ*m) before further use. To initiate the formation of covalent bonds between Gel-MA and HRP-MA, the hydrogels were photo-crosslinked at 395 nm (40 mW cm^−2^) for 1-5 minutes.

The rheological properties of the resulting crosslinked hydrogels were measured with different initial HRP-MA concentrations, starting with 0.625 mg/mL for Gel(HRP) and 1.0 mg/mL for Gel(HRP_1.0_), up to 2.0 mg/mL for Gel(HRP_2.0_). In comparison, we also prepared a gelatin ink without the addition of HRP-MA (Gel). Figure 3A shows the variation of storage modulus (G*’*) and loss modulus (G*’’*) against a change in temperature from 40 ℃ to 10 ℃. The gelation temperature or the gelling point (G*’* = G*’’*) was 21.2 ℃ ± 0.1 in all types of bioinks, indicating that the addition of HRP-MA did not result in any significant effect on the gelling temperature (Figure 3A). When the bioinks were cooled, the G*’* values increased rapidly, overcoming the G*’’* value, and transitioning from a liquid state to a gel-like structure. The yield stress of the hydrogels was also determined by performing shear sweep tests as shown in Figure S4. At 20 °C, all bioinks, regardless of HRP content, had a robust shear-thinning behavior showing a linear decrease of viscosity with an increasing shear rate on a log-log scale, indicating power law behavior, an indicator of good printability (Figure 3B).

**Figure 3.**
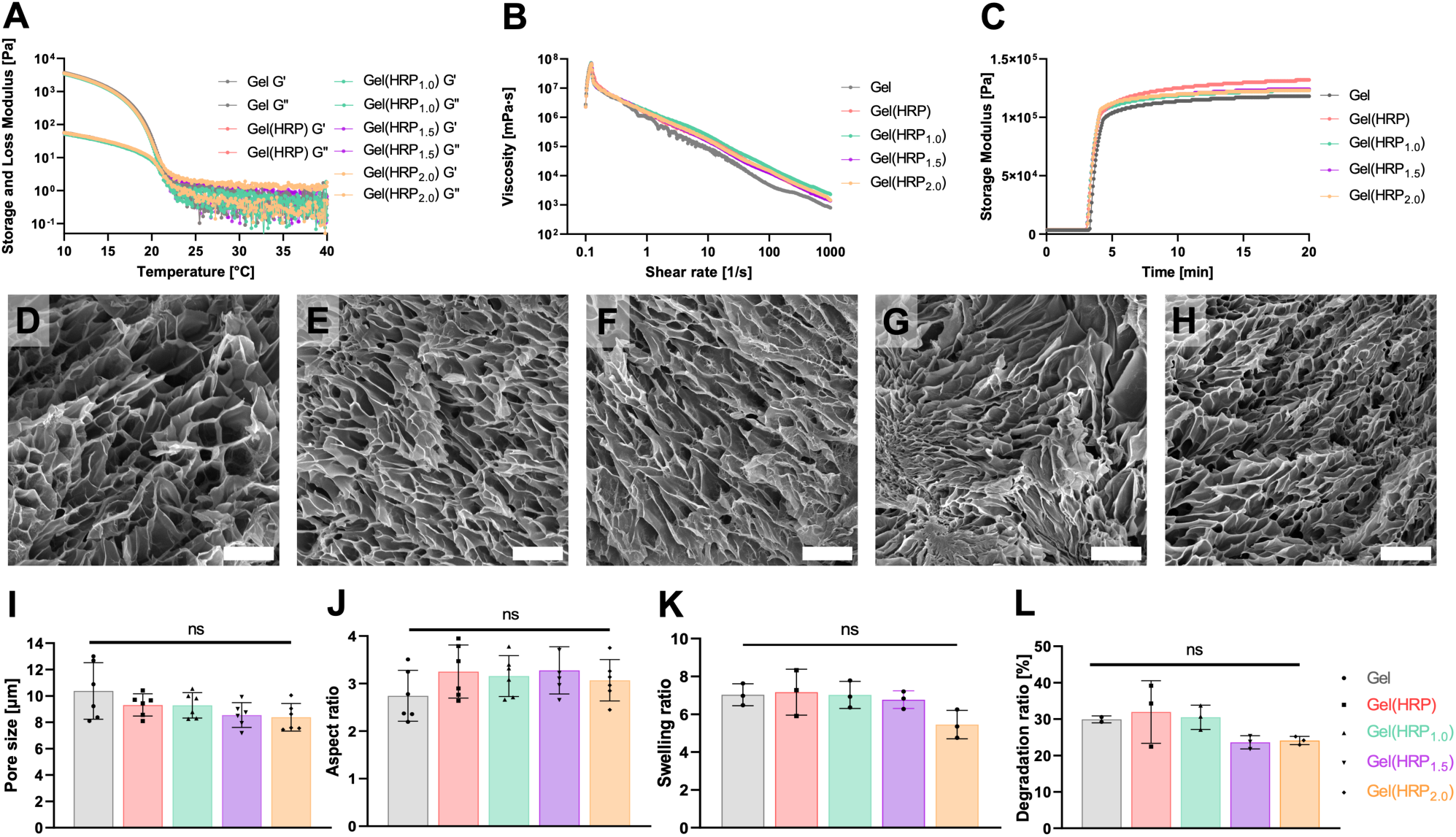
Characterization of hydrogels. (A) Effect of temperature on storage modulus G*’* and loss modulus G*’’*. (B) Viscosity as a function of shear rate. (C) Effect of UV light on the storage modulus G*’*. (D-H) From left to right, scanning electron microscopy images of unmodified Gel-MA ink (Gel), followed by HRP-modified inks Gel(HRP), Gel(HRP_1.0_), Gel(HRP_1.5_), and Gel(HRP_2.0_). Scale bars are 10 *μ*m. (I) Pore size of hydrogels. (J) Aspect ratio of hydrogels (length of pores divided by their width). (K) Swelling ratio of hydrogels after 24 hours in PBS. (L) Degradation ratio of hydrogels after incubation with collagenase I for 48 hours. Error bars represent standard deviation; ns P > 0.05, ANOVA.

To verify the crosslinking of the bioinks, rheological time sweeps were performed while exposing the samples to UV light. Here, G*’* increased rapidly to values above 100 kPa ranging from 118 kPa to 132 kPa after only 60 seconds of UV exposure (395 nm, 40 mW cm^−2^) at 20 °C with no significant differences between individual bioinks (Figure 3C). We also successfully achieved the ability to tune the final stiffness of the hydrogels within a range of 30 kPa to 80 kPa by adjusting the UV light exposure time. Specifically, by varying the UV light exposure time from 5 seconds to 15 seconds, we observed the corresponding stiffness values as depicted in Figure S5.

The structural morphologies of all gel combinations were then analyzed post-freeze-drying with scanning electron microscopy (SEM). All materials presented a microporous structure with uniform pore sizes and relatively smooth inner surfaces. Figure 3D-H indicate that the hydrogel’s pores were elliptical in shape and Gel, Gel(HRP), Gel(HRP_1.0_), Gel(HRP_1.5_), and Gel(HRP_2.0_) had an average pore size (length) of 10.38 *μ*m, 9.31 *μ*m, 9.28 *μ*m, 8.55 *μ*m, and 8.38 *μ*m respectively (Figure 3I). While this trend suggests that increased concentrations of HRP may result in a slight decrease in pore sizes, there were no statistical differences among any samples. The aspect ratio (length of pores divided by their width) was the lowest for Gel (2.73), and all HRP hydrogels had similar values with Gel(HRP), Gel(HRP_1.0_), Gel(HRP_1.5_), and Gel(HRP_2.0_) having aspect ratios of 3.25, 3.16, 3.28, and 3.07 respectively, indicating no statistical difference amongst the different hydrogels (Figure 3J).

Additionally, the swelling and degradation properties of the enzyme bioinks were compared to the unmodified Gel-MA bioinks (Gel). The swelling ratio describes the capacity to absorb liquids and impacts the mechanical and functional properties of hydrogels (Figure 3K). Similarly, the degradation ratio is an important parameter for tissue engineering applications that affect the native cell’s ability to remodel the bioink. Here, the hydrogels were incubated with a collagenase I solution, and the dry weights after 24-hour incubations were compared to the starting weights (Figure 3L).

While there are minor indicators that higher HRP concentrations might affect the crosslinking of the hydrogels and the network strength, the changes are not statistically significant for tangible differences in the hydrogels. Overall, it can be concluded that the addition of HRP in different concentrations has only a minimal effect on the hydrogels’ mechanical integrity.

### 2.3 Enzyme Bioink Activity

To verify that HRP-MA was covalently attached to the Gel-MA scaffold during the photo-crosslinking step, two tests were performed. First, we compared how much functionalized HRP-MA was washed out of bioink droplets compared to native, unmodified enzymes. Then, the remaining enzyme inside each droplet was further analyzed for its catalytic activity and visualized with confocal microscopy.

#### 2.3.1 Enzyme Bioink Washing Test

An ABTS activity assay was run to quantify the amount of unbound enzyme that can be washed out of the bioink (Figure 4). Here, 20 *µ*L UV-treated and crosslinked bioink droplets containing equal amounts of modified and unmodified enzymes were analyzed over time. To mimic physiological conditions, Dulbecco’s Modified Eagle’s Medium (DMEM, high glucose) media was used at 37 °C for 1-hour washes. Unbound enzymes may be retained due to electrostatic interactions with Gel-MA^12, 61^ and physical entanglement in the dense crosslinked polymer network. Therefore, we applied an additional sonication step to help to release loosely entrapped enzymes (Figure 4A).

**Figure 4.**
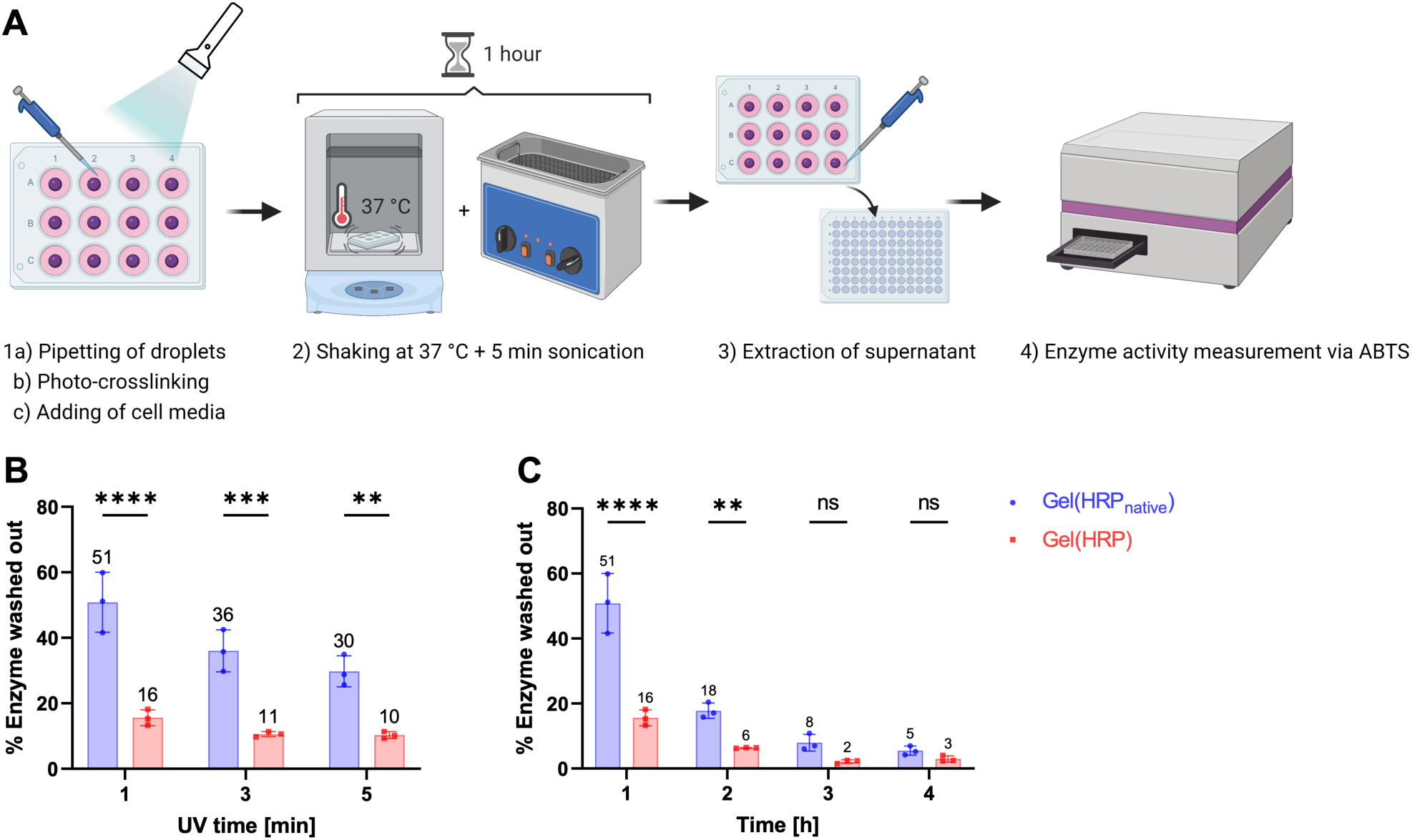
Enzyme bioink washing test to determine crosslinking of the enzyme HRP with the polymer scaffold Gel-MA. (A) Schematic experiment set-up of one washing cycle. (B) Enzyme amount washed out of a 20 *µ*L droplet measured via ABTS assay of 10 *µ*L washing solution after 1 hour in % of the total enzyme amount, UV (395 nm) times of 1 min, 3 min, and 5 min were applied. (C) Enzyme amount washed out of a 20 *µ*L droplet (1 min of UV crosslinking) measured via ABTS assay of 10 *µ*L washing solution in dependency of time (1-4 washing cycles/hours) in % of the total enzyme amount. *P < 0.05, **P < 0.01, and ***P < 0.001, two-way ANOVA.

The total initial amount of enzyme within each droplet was determined by dissolving equal droplet volumes without crosslinking in media. An ABTS activity test revealed that both Gel(HRP) and Gel(HRP_native_) droplets contain the same initial enzyme amount and activity (Figure S6). Additionally, the impact of the photocrosslinking process on the enzyme activity was evaluated after the degradation of the bioink droplets by collagenase I and quantifying the activity of the released enzymes. The results show that the enzyme activity was unharmed by the droplet formation and the photo-crosslinking step (Figure S7). In summary, this confirms, that all initial droplets contain the same amount of enzymes, and that their native activity is not affected by the covalent attachment to the polymers.

For the enzyme washing tests, the amount of released unbound enzyme was measured after each washing cycle by performing activity tests. After 1 hour of shaking in media, around 51 ± 8 % of HRP_native_ was washed out of the bionk droplets and is detectable in the surrounding solution (Figure 4B). In comparison, the functionalized HRP-MA was retained more effectively inside the gel and only 16 ± 3% of enzymes were detected in the washing solution. As expected, longer UV crosslinking times (3 min, 5 min), increase the Gel-MA network density and restrict the overall release of both types of enzymes. Comparing the relative confinement of functionalized HRP-MA within the bioink shows a 35 % higher retention with 1 min of UV crosslinking (25 % with 3 min, and 20 % with 5 min). For all further studies, we selected a 1 min UV crosslinking time since we observed a high crosslinking efficiency, while at the same time, having a relatively low network density for good nutrient diffusion and cell growth.

The bioink droplets treated with 1 min of UV light were further analyzed every hour for the next 3 hours (Figure 4C). Between each subsequent time point, less and less enzyme was found to be released from either hydrogel. In all cases, more HRP_native_ is released compared to HRP-MA. After 4 hours, the detected amount of enzyme in the surrounding washing solution reaches a lower limit of around 5 and 3 %, respectively. This results in a cumulative release of 82 % of HRP_native_ and only 27 % of HRP-MA. The low levels of HRP-MA release are most likely caused by a slightly reduced conjugation efficiency due to oxygen inhibition of the crosslinking reaction in the hydrogels. At the same time, the reason why not all unbound enzymes are immediately washed out of the gels is an additional electrostatic interaction between the large biomolecules and the polymer network. Overall, the washing test results highlight the importance of a covalent enzyme conjugation compared to the addition of native enzymes which are almost completely depleted after just a few hours.

We expect to see an even more significant relative retention effect of HRP-MA over prolonged time, increased salt concentrations, changing pH, as well as due to the incorporation of cells. Under these challenging conditions, covalent conjugation should prove even more beneficial, and weakened non-covalent interactions or looser network structures will further accelerate the washing out of unbound enzymes. For example, an ABTS assay conducted on cell-laden hydrogels revealed that those containing HRP-MA retained the enzyme for a full 7 days and maintained its activity. In contrast, nearly all of the HRP_native_ was depleted under standard physiological cell culture conditions (see Figure S8). This demonstrates that the enzymes are not only effectively crosslinked and retained within the hydrogels but also remain fully functional.

#### 2.3.2 Enzyme Bioink Activity Test Inside the Gel

For a direct activity test of enzymes inside of the hydrogels, bioink droplets were exposed to a minimum of 5 washing cycles (including one overnight cycle) and the remaining enzyme amount inside the gel was analyzed using a fluorescence method to visualize the enzyme activity via confocal microscopy inside the gels. Here, the substrate Amplex Red is oxidized to fluorescent resorufin in the presence of the HRP (Figure 5A).^55, 62^ Gel-MA (Gel) droplets without HRP served as controls.

**Figure 5:**
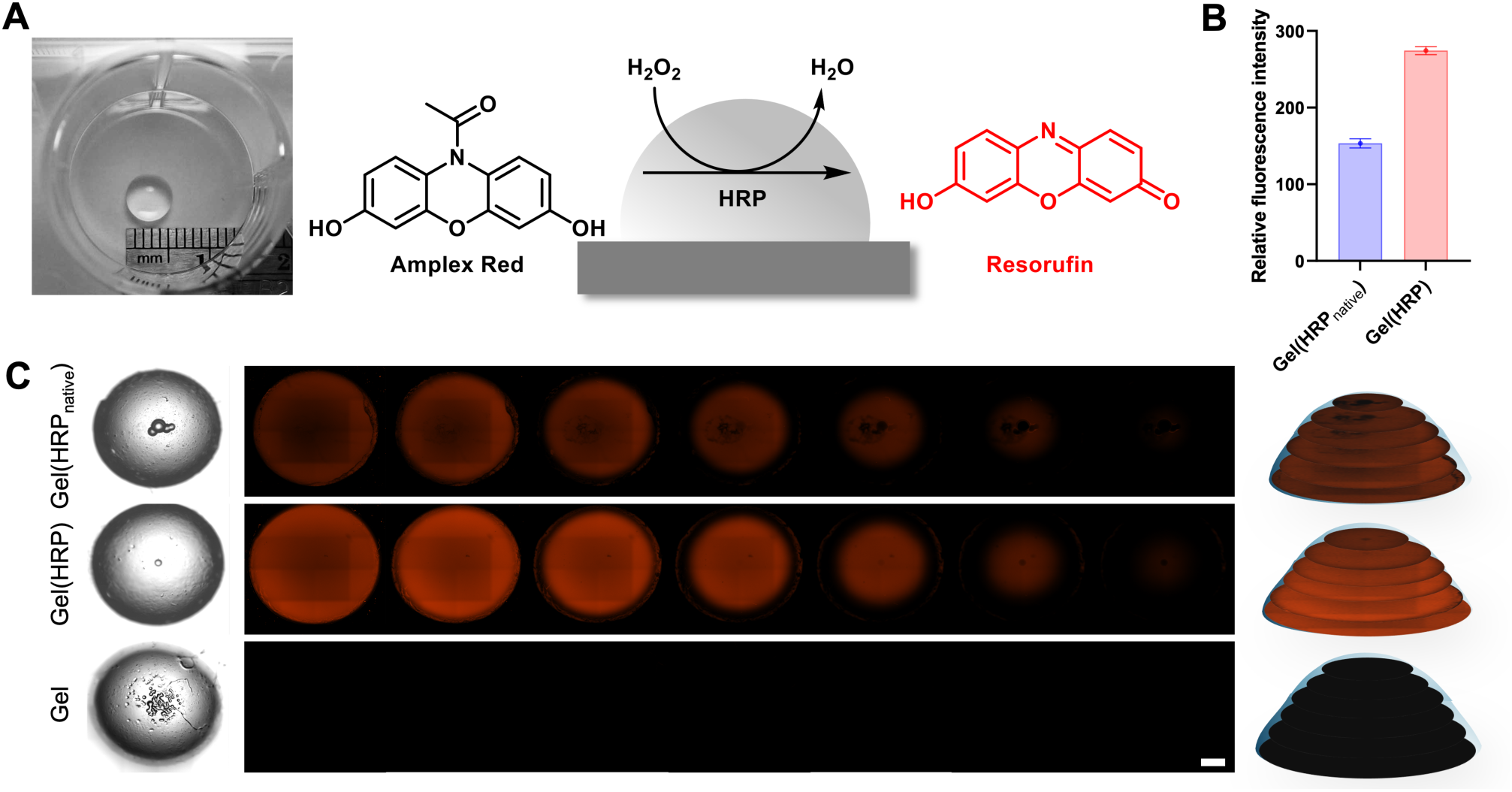
Enzyme bioink activity test inside the gel to determine crosslinking of the enzyme HRP with the polymer scaffold. (A) Image of a bioink droplet; schematic reaction of Amplex Red oxidation to resorufin in the presence of HRP. (B) ImageJ analysis of mean fluorescence in each layer of the droplets of Gel(HRP_native_) and Gel(HRP); Error bars represent standard deviation. (C) Left, brightfield images of Gel(HRP_native_), Gel(HRP) and Gel (objective 5x, NA = 0.35); Middle, confocal images of z-stack plane images of enzyme bioink droplets after washing treated with Amplex Red and H_2_O_2_ taken on Zeiss CD7 LSM 900 (Ex/Em = 568/581 nm); Scale bar = 1 mm; Right, z-stack plane images from C reassembled into 3D droplets.

For this, confocal z-stack images comprising 7 slices capturing several layers of the full droplet were measured (Figure 5C). A high fluorescence intensity of resorufin indicates a high enzyme activity and hence a large amount of enzyme, respectively. As expected, the enzyme-free droplet (Gel) did not display any fluorescence, whereas the enzyme-containing droplets Gel(HRP_native_) and Gel(HRP) showed fluorescence, with Gel(HRP) displaying a higher intensity than the native form. The uniformity of the fluorescence signal shows a near homogeneous HRP distribution throughout the hydrogel, with slightly restricted penetration of the assay substrates to the droplet center. The enzymatic activity of each droplet was assessed by calculating the mean fluorescence intensity of each z-slice throughout the full droplet with ImageJ (Figure 5B). Droplets with covalently attached enzymes have a significantly higher fluorescence and thus enzyme activity. Image analysis showed that almost twice as much HRP-MA is retained compared to native HRP, which is washed out more readily.

The in-droplet activity results showed a similar trend as the washing tests earlier. The slightly smaller difference in enzyme retention can most likely be explained by the aspect that the bioink washing test favors looking at enzymes closer to the droplet surface, whereas the Amplex Red assay can analyze the center of a droplet. Non-attached enzymes deeper within the gel are more effectively retained due to stronger physical entrapment, in contrast to enzymes near the surface, which are more prone to being washed out.

By printing smaller-sized bioink droplets, we also showed that the size of the structure has minimal effect on enzyme retention. Lastly, we were also able to demonstrate the tunability of our approach by conjugating 50% of the initial enzyme amount to a droplet, consequently observing half of the corresponding activity (Figure S9).

#### 2.3.3 Enzyme Activity after 3D Printing

Extrusion-based printing has the benefit of positioning the catalytic enzyme hydrogel precisely at targeted locations and has the potential in further studies to form complex functional 3-dimensional structures. As proof-of-principle and to explore the general printability, the enzyme bioink was placed in simple three-dimensional shapes and its enzyme activity was captured via Amplex red oxidation. For this, a grid structure, the outline of the country of Australia, a star, and a round vessel were printed with a modified Lulzbot Mini 2 3D printer and captured via confocal microscopy (Figure 6).

**Figure 6.**
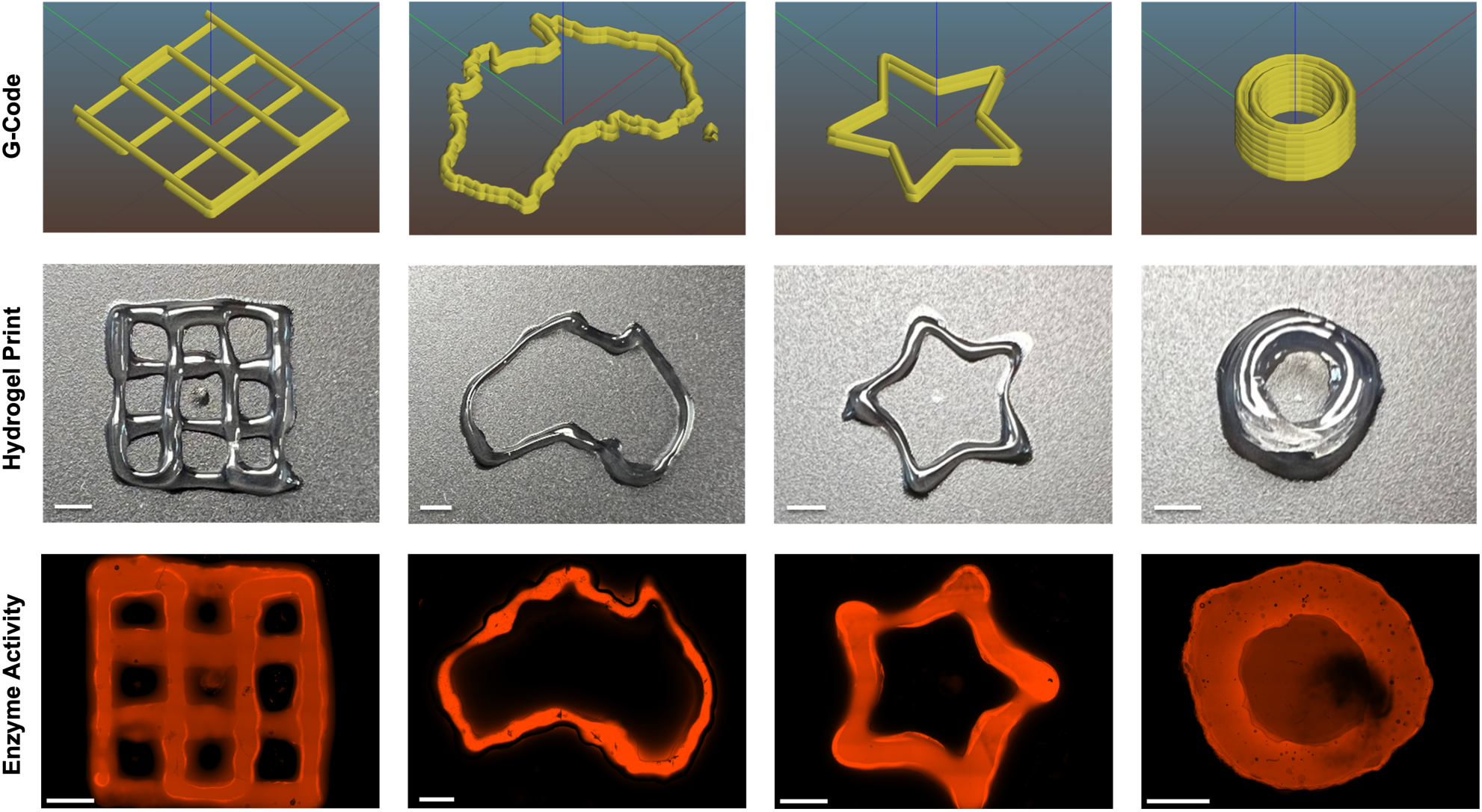
3D Extrusion-based bioprinting with an enzyme-containing Gel(HRP) ink; 3D structures from left to right: grid, Australia, star, vessel; Top: 3D models by Slic3r software; Middle: Photos of the printed structures; Bottom: confocal images of materials treated with Amplex Red and H_2_O_2_ (Zeiss CD7 LSM 900). Scale bar 2 mm.

In the case of the circular vessel, we explored the application as a basic reaction vessel. For this, the cavity inside of the vessel was filled with an Amplex Red solution and H_2_O_2_. The immobilized HRP in the vessel walls oxidized the substrate to resorufin, which penetrated the hydrogel walls and stained them red. These experiments demonstrate the printability of the enzyme bioink with a basic extrusion bioprinter and exemplify that the enzyme activity was preserved in the printing process.

### 2.4 Cytocompatibility of Adipose-Derived Stem Cells (ADSC) in Enzyme Bioink Constructs

To evaluate the general cytocompatibility of HRP-MA modified bioinks, we performed cell viability tests on ADSC-laden hydrogels. The hydrogels without enzymes (Gel) and crosslinked with enzymes (Gel(HRP)) were compared after being placed and cultured in square plastic molds. In addition, we also analyzed the cell viability of extrusion-printed hydrogels (Gel(HRP) 3D-printed). For days 1 and 7, confocal images show high levels of live cells (green) with minimal dead cells (red) for all gel conditions (Figure 7A). Quantification revealed cell viability above 90% for all conditions across all time points (Figure 7B). This is notable since high stiffness can affect the cell viability of ADSCs. For Day 7, the hydrogel without enzymes (Gel) had a viability of 93.5 ± 3% while the HRP-MA containing hydrogels had comparable cell viability. There was no statistical difference between the control and HRP-MA conditions (P > 0.999), indicating that the inclusion of HRP-MA does not affect cell viability. In addition, confocal imaging at day 7 indicates better cell spreading in HRP-MA hydrogels, particularly in the 3D-printed form. Here, the cells also show clear signs of proliferation with elongated cell structures (Figure 7A). When counting live cells per image, a significantly increased number of cells (P < 0.001) can be observed in Gel(HRP) 3D-printed, whereas the control (Gel) shows only a slight increase in cells from day 1 to day 7 (Figure 7B).

**Figure 7.**
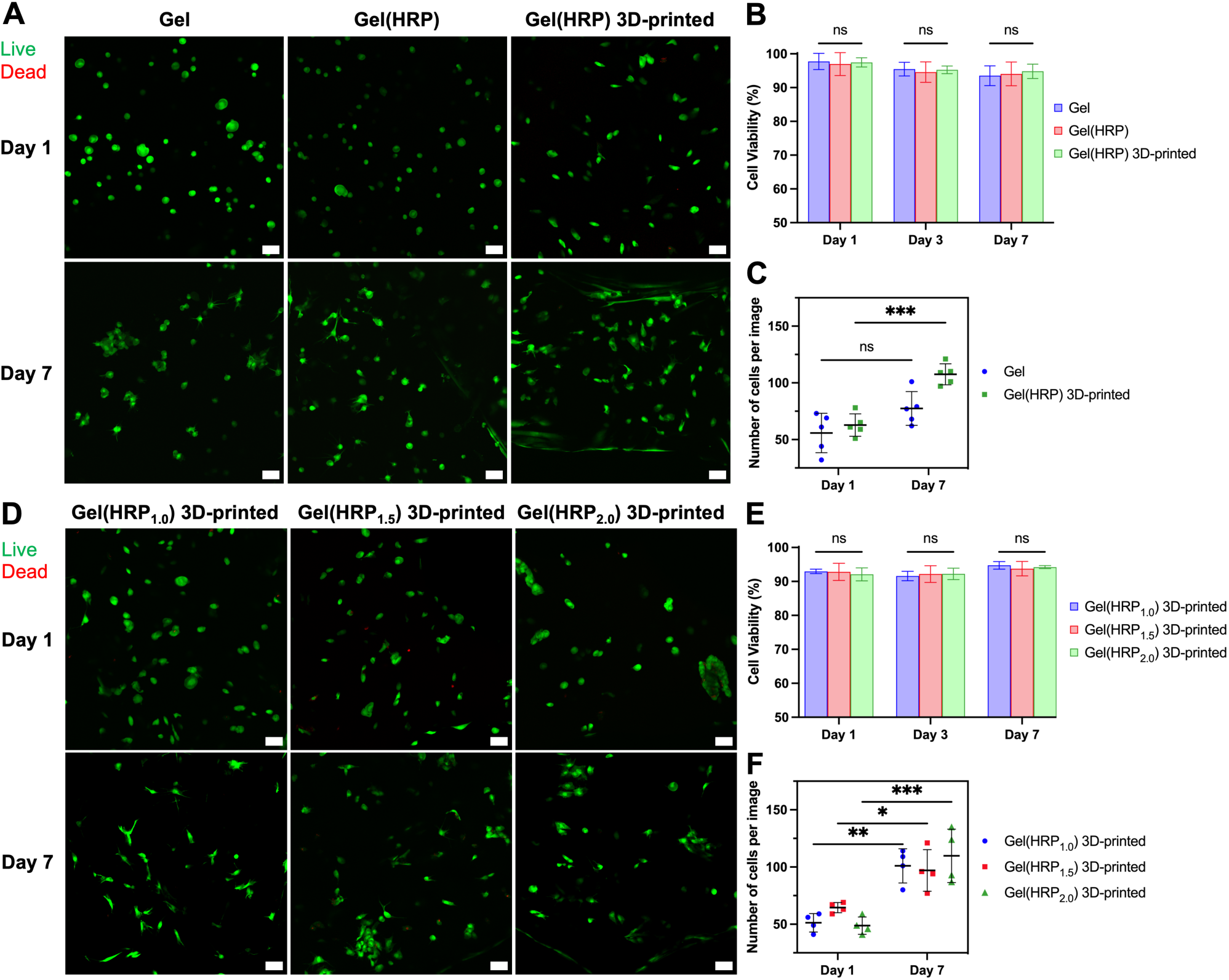
(A) Live (green, calcein AM) dead (red, ethidium homodimer 1) stain of ADSCs-loaded in Gel, Gel(HRP), and Gel(HRP) 3D-printed hydrogel. (B) Quantification of cell viability of ADSCs over 7 days. (C) Quantification of the average number of cells per image (n = 5, number of images). (D) Live (green, calcein AM) dead (red, ethidium homodimer 1) stain of ADSCs loaded in Gel(HRP_1.0_) 3D-printed, Gel(HRP_1.5_) 3D-printed, and Gel(HRP_2.0_) 3D-printed hydrogels. (E) Quantification of cell viability of ADSCs. (F) Quantification of the average number of cells per image (n = 4, number of images). Scale bar = 50 *μ*m. *P < 0.05, **P < 0.01, and ***P < 0.001, two-way ANOVA.

To further assess the cytocompatibility of the enzyme inks, we similarly performed live-dead tests on ADSCs in Gel-MA hydrogels with higher concentrations of HRP-MA (1.0–2.0 mg/mL) (Figure 7D). Again, cell viability for all concentrations of HRP-MA was more than 90% for all time points (Figure 7E). In addition, we also assessed the average cell area within the hydrogels as an indicator for improved cell viability (Figure S10). Overall, the results indicate that the HRP-containing gels show high cell viability, and the inclusion of enzymes does not interfere with the general cytocompatibility of gelatin bioinks.

## 3. Conclusion

We demonstrate a universal approach for the preparation of functional and catalytical active bioinks. Using HRP as a model enzyme, we describe a conjugation approach to covalent link proteins to standard Gel-MA bioinks and therefore increase their retention and bioavailability. A mild chemical modification introduces multiple methacrylate groups on the surface of the enzymes, without denaturing the three-dimensional structure or affecting the biocatalytic activity. This allows a photo-crosslinking of gelatin with free methacrylate groups after printing. The tethering of the enzyme to the bioink scaffold significantly decreases their premature loss from the hydrogel and results in a high sustained catalytic activity.

We demonstrate the cytocompatibility of the enzyme bioink and its printing capabilities towards applications in wound healing, regenerative medicine, and bioactive coatings. We envision that this approach can be universally applied to other enzymes enabling a wide array of biocatalytic transformations. Examples include ROS scavenging, antibacterial activity, or oxygen generation, offering the possibility to incorporate diverse biochemical activities into biofabricated matrices.

## 4. Experimental Section

### Materials

Gelatin (300 bloom, Type A), methacrylic anhydride (MAA), 2,2’-azino-bis(3-ethylbenzothiazoline-6-sulfonic acid) (ABTS), fluorescamine, deuterium oxide (D_2_O), 2,2’-Azino-bis(3-ethylbenzothiazoline-6-sulfonic acid) diammonium salt (ABTS), horseradish peroxidase (Type II, 150-200 U/mg), Amplex Red, FGF2, and DMSO (anhydrous) were purchased from Sigma-Aldrich; Dulbecco′s Modified Eagle′s Medium (high glucose, pyruvate, no glutamine, DMEM), collagenase (type I) was purchased from Thermo-Fischer; Live-dead kit (calcein and ethidium homodimer) was purchased from Life Technologies Australia; lithium phenyl-2,4,6-trimethylbenzoylphosphinate (LAP) was purchased from Ambeed; hydrogen peroxide (27-30%) and dimethyl sulfoxide (DMSO) was purchased from Chem-Supply Pty Ltd.

### Instrumentation

^1^H-NMR spectra were recorded using a 300 MHz Bruker Topspin Fourier spectrometer. Infrared (IR) spectra were recorded using a Nicolet Avatar 330-IR (ATR-unit) spectrometer. Absorption and fluorescence measurements were performed on a Synergy H4 (BioTek) plate reader. Confocal microscopy images were captured using a Zeiss LSM 900 CD7 and Zeiss LSM 800 microscope. Applied Photophysics Chirascan Plus CD was used for CD spectroscopy. Anton Paar MCR 302 rheometer with a parallel plate geometry (25 mm disk, 1 mm measuring distance, 600 *μ*L of liquid sample) was used for rheological tests. Furthermore, the ultrasonic bath Thermoline Scientific Power sonic 410, the multi centrifuge OHAUS Frontier FC5706 230V and the freeze-dryer Christ Alpha 1-2 LDplus were used.

### Methacrylation of Gelatin (Gel-MA)

Methacrylate-modified gelatin (Gel-MA) was prepared according to the previously described one-pot method by *Shirahama*.^52, 53^ In brief, gelatin was dissolved in 10 w/v% sodium carbonate– bicarbonate (CB) buffer (0.25 M, from Sigma Aldrich, pH was adjusted to 9) at 50 °C. MAA (94%, 0.1 mL) was added slowly to a solution of 1g gelatin under vigorous stirring. After stirring the mixture at 50 °C for one hour, the pH was readjusted to 7.4 with 6 M hydrochloric acid to stop the reaction. The solution was then purified by filtration and dialysis (MWCO 14 kDa, Thermo Fisher Australia) against deionized water at 50 °C for 3 days before freeze-dried for 4 days and stored at −20 °C until further use. The product is a colorless foam-like material (yield 83%). ^1^H-NMR (Figure S1) and FTIR (Figure S3) were carried out to verify the amine substitution, and the degree of methacrylation was quantified via fluorescamine assay.

### Methacrylation of Horseradish Peroxidase (HRP-MA)

For the methacrylation of horseradish peroxidase, the procedure by *Ferracci et al*.^39^ and *Smith et al.*^49^ was adapted in parts. Here, 1 mg (22.73 nmol) of HRP was dissolved in 1 mL of CB buffer (0.25 M, pH 9) and 1 *µ*L (0.1 w/v %) of a stock solution of MAA in DMSO (86.43 *µ*L in 413.57 *µ*L) was added to result in an MAA/lysine ratio of 8:1 (1.09 *µ*mol MAA, 0.168 mg, 0.173 *µ*L of a 94% MAA solution). The resulting mixture was stirred for 8 h at room temperature. The product was purified four times by spin filtration in Amicon® Ultra 15 mL filter devices (MWCO 30 kDa) from Sigma-Merck Millipore at 6000 rpm for 5 min at room temperature and then lyophilized overnight. The product is a light brown foam-like material (yield 86%).

### Determination of Methacrylation via Fluorescamine Assay

All samples (HRP, HRP-MA) were adjusted to a concentration of 1.6 mg/mL via absorbance measurement at 280 nm. HRP was used as an external standard in a concentration range of 0.016 mg/mL to 2 mg/mL. For the assay, 125 *µ*L PBS (0.01 M, pH 7.4), as well as 25 *µ*L sample/standard, was added in triplicates into each well of a 96-black-well-microplate (flat bottom). Just before the measurement, 50 *µ*L of 0.3 mg/mL freshly prepared fluorescamine solution in DMSO was mixed in before analyzed with a fluorescence plate reader (Ex/Em = 380/460 nm); The degree of substitution (DS) of amines to methacrylate groups was calculated as follows:

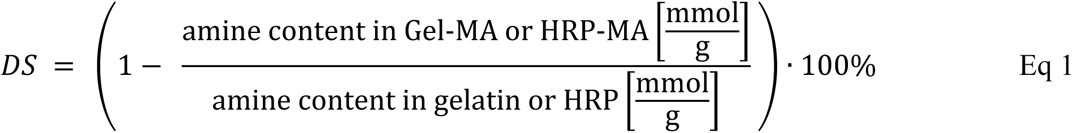

### CD Spectroscopy

CD spectra were recorded at 20 °C with a total enzyme concentration of 0.1 mg/mL in CD buffer (containing 10 mM potassium phosphate and 50 mM Na_2_SO_4_) using quartz cells with a path length of 1 mm. Spectra were corrected by subtraction from the background (buffer). Data points were collected at a resolution of 1 nm. CD spectra were analyzed using the software Pro-Data Viewer 4.7.0.194.

### Determination of Enzyme Activity via ABTS Assay

The peroxidase activity of HRP and HRP-MA was measured by using 2,2’-azinobis-(2-ethylbenzthiazoline-6-sulfonate) (ABTS, 0.25 mM) following a slightly modified protocol.^63^ Briefly, ABTS was dissolved at a concentration of 2 mM in 10 mM phosphate buffer pH 6. The enzymes HRP and HRP-MA were dissolved at a concentration of 40 *µ*M and their concentration was adjusted to the same values by absorption measurements at 280 nm. The H_2_O_2_ concentration ranged between 0.4 mM and 30 mM (in ddH_2_O). In a 96-clear well plate, 100 *µ*L ABTS solution and 25 *µ*L enzyme sample were added. 50 *µ*L of the respective H_2_O_2_ concentrations were added to the mixture and the absorption was measured at 410 nm for 3 min every 10 s in a kinetic cycle. All measurements were carried out in triplicate. Data was analyzed using GraphPad Prism 9.

### Enzyme-functionalized Bioink

In a 4 mL glass vial equipped with a magnetic stir bar, an amount of 55 mg Gel-MA (11 w/v %) in 418.74 *µ*L of 1x PBS was put into a water bath at 40 °C for dissolution. 50 *µ*L of a LAP solution (0.25 w/v %, stock solution: 25 mg/ml in ddH_2_O) and 31.25 *µ*L enzyme (0.625 mg/mL, stock solution: 10 mg/mL) were added and the mixture was filtered once via syringe filter (Ø 45 *µ*m) before use. Unless for the rheology experiments which examine the bioink state in its non-crosslinked form (shear-thinning and viscosity properties), the hydrogels were always further photo-crosslinked at 395 nm using a UV light torch (100 LED 395 nm UV ultraviolet flashlight blacklight torch) at a power of 40 mW cm^−2^ for 1, 3 or 5 minutes.

For the verification of the crosslinking, the functionalized enzyme hydrogel Gel(HRP) was compared with the hydrogel that contains only native enzyme mixed-in (Gel(HRP_native_)). For these experiments, the concentrations were equalized via absorption measurement at 280 nm. Additional concentrations (Gel(HRP_1.0_) 1.0 mg/mL, Gel(HRP_1.5_) 1.5 mg/mL, Gel(HRP_2.0_) 2.0 mg/mL) were used to investigate the impact of the functionalized enzyme on the hydrogel material properties such as rheological, swelling, degradation, and cytocompatibility.

### Rheology of Bioink

All the rheological measurements on Gel, Gel(HRP), Gel(HRP_1.0_), Gel(HRP_1.5_), and Gel(HRP_2.0_) hydrogels were performed on an Anton Paar MCR 302 rheometer with a parallel plate geometry (25 mm disk, 1 mm gap size, 600 *μ*L of liquid hydrogel). All the hydrogel samples were placed on the plate at 40 ℃ to fill the gap between the plates. The temperature gelation measurement was evaluated by measuring storage modulus (G*’*) and loss modulus (G*’’*) as a function of temperature ramping from 40 ℃ to 10 ℃ at the rate of 1 ℃/min, at a constant frequency of 1 Hz and a constant strain of 0.2 %. The measurements of viscosity and shear stress were performed by varying the shear rate from 0.1 to 1000 s^−1^ to give us the shear thinning behavior of the hydrogel samples. For UV crosslinking measurements, the storage modulus of the samples was measured for 20 min and a UV lamp (with 395 nm UV light at 40 mW cm-^2^) was placed underneath to illuminate the samples after 4 minutes with UV light for 1 min through quartz crystal stage.

### SEM Imaging

The porous structures of Gel and different Gel(HRP) hydrogels were analyzed using an FEI Nova NanoSEM 450. The hydrogels were frozen and subsequently freeze-dried. After the samples were completely dry, they were cut, and their cross sections were coated with 5 nm chromium and 10 nm platinum layers using Leica ACE600 and EMITECH K575X sputter coaters respectively. SEM images were captured at a high voltage of 5 kV using secondary electron mode.

### Swelling Ratio

50 *µ*L droplets of bioink were photo-crosslinked and submerged with 3 mL of PBS (1x, pH 7.4) in Petri dishes (Sarstedt, Ø 45mm). After shaking at 80 rpm and 37 °C in an orbital shaker for 24 hours, the samples were removed, lightly blotted dry with a lint-free wipe, and weighed (W_(swell)_). The swollen samples were then lyophilized and weighed again (W_(dry)_). To determine the initial dry weight (W_(dry_,_i)_), droplets were lyophilized and weighed immediately after the initial preparation and photo-crosslinking (without PBS treatment). The percentage mass swelling ratio was calculated as follows:

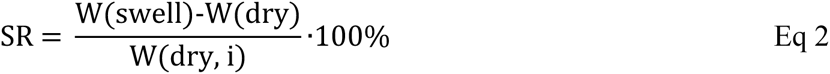

### Degradation Ratio

For the degradation test, 90 *µ*L of warm liquid bioink in 1.5 mL Eppendorf tubes were photo-crosslinked. The gels were then immersed with 0.5 mL PBS containing 2 U mL^−1^ collagenase type I solution at 37 °C in an orbital shaker at a shaking speed of 80 rpm for 24 hours. The collagenase solution was discarded, and the tube was washed once with ddH_2_O water. Excess water was removed, and 0.5 mL of fresh collagenase solution was added. After another 24 hours, the samples were lyophilized to determine the dry weight (W(t)). With the initial lyophilized weight (*W*(0) = dry weight before soaking) of the bioink gels, which was measured with separate samples, the degradation ratio could be calculated as follows:

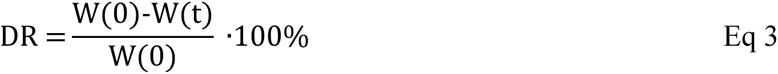

### Enzyme Bioink Washing Test

The enzyme bioink containing either the native HRP (Gel(HRP_native_)) or the methacrylated HRP (Gel(HRP)) was pipetted in 20 *µ*L droplets at 37 °C into separate wells of a 12-well cell culture plate (CoStar, 3513). Subsequently, these droplets were photo-crosslinked at 395 nm for 1, 3 or 5 min, and immersed with 2 mL DMEM. As a control, 20 *µ*L of enzyme bioink was pipetted directly into a 2 mL media, mixed, and exposed to an equal amount of UV light (395 nm) as the droplet of shaking at 80 rpm and 37 °C well as 5 min of sonication, 10 *µ*L supernatant media was extracted and its enzyme activity was measured via ABTS assay. Therefore, 40 *µ*L ddH_2_O, 100 *µ*L ABTS solution, and 50 *µ*L of H_2_O_2_ (stock solution 0.1 mM in ddH_2_O) were added additionally into a 96-well plate (Sarstaedt) and the absorbance was measured immediately at 410 nm for 3 min every 10 s in a kinetic cycle. The media was discarded, residues removed, and 2 mL of fresh media was added. After another hour in the incubator and 5 min of sonication, the supernatant was measured for enzyme activity again to monitor the enzyme washed out over time. The measured values were analyzed via GraphPad Prism 9.

### Enzyme Bioink Activity Test Inside the Gel

After five washing cycles, the activity inside the gels was visualized via confocal microscopy through the oxidative conversion of Amplex Red to resorufin. Tile images of different bioink droplets (Gel, Gel(HRP) and Gel(HRP_native_)) were visualized in brightfield and confocal mode (objective 5x, NA 0.35; Ex/Em = 568/581 nm). Tile images of identical size and z-height from the plate bottom are comprising 10 slices of z-stack covering a full droplet.

Before imaging, media was removed, and the droplets were rinsed once with ddH_2_O. Afterward, 2 mL of PBS (0.01 M, pH 7.4) was added to each of the wells containing the bioink droplets. Then, 10 *µ*L of Amplex Red solution (stock solution 10 mM in DMSO) was added, mixed, and incubated for 5 min before adding 10 *µ*L of H_2_O_2_ (stock solution 0.1 mM in ddH_2_O) and incubated for additional 10 minutes. The solution was discarded, and any excess was removed gently with lint-free wipes. Image analysis was performed with Imaris software (version 9.9.0). The fluorescence intensity was calculated with ImageJ by selecting the droplet area of each plane and carrying out a mean measurement which was then summed up to a total value.

### Printing of Enzyme Bioink

The 3D bioprinter was modified from the commercial Lulzbot Mini 2 3D printer by replacing the preinstalled printhead with a screw extrusion syringe head. All bioprinting steps were conducted at 21 °C except for the cell studies, which were performed at 19 °C due to the preset local room temperature. The bioprinting speed was kept constant at 1.66 mm/s. Printing GCode was generated with help of the slicing software Slic3r, and the extrusion rate was standardized via Python to obtain a constant extrusion per distance value of 0.06 throughout the full 3D structure. Additionally, the GCode was modified in order to avoid a printing delay at the start and to a pull-up of the printed structure at the end.

After preparing the hydrogel mixtures at 40 °C, the bioink-containing syringe was cooled down to room temperature before being extruded by the bioprinter equipped with a 21G Nordson EFD precision needle. Immediately after printing into separate wells of a 12-well cell culture plate, the structures were further UV-photo-crosslinked at 395 nm for 1 min at a power of 40 mW cm^−2^. Afterward, 2 mL of PBS (0.01 M, pH 7.4) was added to each of the wells containing the bioink structures. Then, 10 *µ*L of Amplex Red solution (stock solution 10 mM in DMSO) was added, mixed, and incubated for 5 min before adding 10 *µ*L of H_2_O_2_ (stock solution 0.1 mM in ddH_2_O) and incubated for additional 10 minutes before the solution was discarded and any excess was removed gently with a lint-free wipe. For the vessel structure, 0.5 mL of 1x PBS was inserted into the center of the vessel structure. Then, 2 *µ*L Amplex Red solution (10 mM in DMSO) was added and incubated for 5 min followed by the addition of 2 *µ*L of H_2_O_2_ (0.1 mM in ddH_2_O) with a 10 min incubation time. The vessel was imaged while still having the solutions present. Image analysis was performed with Imaris software (version 9.9.0).

### Cell Culture Methods

Adipose-derived stem cells (ADSCs) were cultured in low glucose DMEM supplemented with 10 vol% fetal bovine serum (FBS) and 1 vol% penicillin/streptomycin and 0.01 vol% FGF2 at 37 ℃, 5% CO_2_ in a humidified incubator. The media was changed every 48 hours, and cells were passaged at 85-90% confluency. All cells used in this work were between passages 5–11.

### Preparation of Cell-laden Hydrogels

To prepare cell-laden hydrogel hydrogels, 11 w/v% concentration of Gel-MA (DS 98.6%) with 2.5 mg/mL LAP was used in all samples and dissolved in 1x PBS at 40 ℃. The enzyme-modified hydrogels (Gel(HRP)), contained in addition 0.625 mg/mL HRP-MA (DS 86%). After complete dissolution, ADSCs (1.5 million/mL) were loaded into the hydrogel solutions. Subsequently, 80 *μ*L of gels were placed into 6⨯6⨯2.5 mm plastic molds and crosslinked using a 395 nm UV light torch (eBay; 100 LED 395 nm UV Ultraviolet Flashlight Blacklight Torch) at 40 mW cm^−2^ for 1 minute. For 3D-printed hydrogels, Gel(HRP) with 1.5 million ADSCs per mL were loaded into a 1 mL Livingstone syringe and kept at room temperature for 15 minutes. The syringe was then loaded into the 3D bioprinter to print grid-like structures with a 21G Nordson general-purpose tip. The grids were then crosslinked using the UV torch light for 1 min. All cell-laden hydrogels were placed in well plates and cultured for 7 days with 1 mL media. Media changes were made during days 1, 3 and 5. Hydrogels with higher concentrations of HRP-MA were 3D-printed to evaluate the toxicity of HRP-MA in high concentrations. These three different concentrations of HRP-MA (1.0, 1.5, and 2.0 mg/mL) were mixed with Gel-MA, corresponding as Gel(HRP_1.0_) 3D-printed, Gel(HRP_1.5_) 3D-printed, and Gel(HRP_2.0_) 3D-printed.

### Cell Viability of Incorporated ADSCs in Hydrogels Using Live-Dead Staining

On days 1, 3 and 7 after the preparation of cell-incorporated hydrogels, the viability of ADSCs within the hydrogels was investigated using a live-dead staining assay. Here, media was removed, and gels were washed once with 1x PBS. Then 500 *μ*L of 1x PBS with calcein-AM (2 *μ*M) and ethidium homodimer-1 (4 *μ*M) was added, and the gels were incubated at 37 ℃ for 45 minutes before washing two times with 1x PBS for 5 min each. The gels were then imaged immediately on a Zeiss LSM 800 confocal microscope (ZLSM) by taking z-stack scans (10x objective, 10 slices over 18 *μ*m). The images were analyzed using Fiji ImageJ software. For every hydrogel, 3 replicates and from each replicate, 2 images with multiple z-stacks from different parts of the gel were analyzed. First, the individual z-stack images of calcein-AM (green) channel (Ex/Em = 488/520 nm) were projected or stacked together using the maximum intensity option in the projection type. The images were then thresholded in the calcein-AM channel to outline the cells, which was followed by watershed segmentation of the image to cut apart connected cell components into separate ones. Individual cells were analyzed using 0.25-1.00 circularity and counted. Similarly, the steps were followed for the ethidium homodimer-1 channel (Ex/Em = 528/617 nm) and dead cells were counted. The viability was calculated by analyzing 5 images per sample and taking the percentage of live to dead cells.

### Statistical Analysis

Statistical analyses were performed using GraphPad Prism (version 9) software while all experiments were carried out in triplicate and the data were analyzed using a two-way ANOVA with Tukey’s Post Hoc HSD analysis unless stated otherwise. Differences were considered significant when P < 0.05. Significance was indicated by * P ≤ 0.05; ** P ≤ 0.01; *** P ≤ 0.001, **** P ≤ 0.0001.

## Author Contributions

The manuscript was written through the contributions of all authors. All authors have approved the final version of the manuscript and declare no competing financial interest.

**Luca A. Altevogt**: Conceptualization, Methodology, Formal analysis, Investigation, Data curation, Writing - Original Draft, Writing - Review & Editing, Visualization

**Rakib H. Sheikh**: Conceptualization, Methodology, Formal analysis, Investigation, Data curation, Writing - Review & Editing, Visualization

**Thomas G. Molley**: Methodology, Investigation, Writing - Review & Editing

**Joel Yong**: Investigation, Writing - Review & Editing

**Kang Liang**: Writing - Review & Editing, Supervision

**Patrick Spicer**: Writing - Review & Editing, Supervision

**Kristopher A. Kilian**: Writing - Review & Editing, Supervision

**Peter R. Wich**: Conceptualization, Resources, Writing - Original Draft, Writing - Review & Editing, Supervision, Project administration, Funding acquisition

## Acknowledgments

The authors acknowledge the support of the School of Chemical Engineering and the Faculty of Engineering at the University of New South Wales, Sydney (UNSW, Sydney), as well as the support of the Australian Centre for NanoMedicine (ACN). L.A. is supported by a Scientia PhD Fellowship (UNSW, Sydney). R.H.S. is supported by an Australian Government Research Training Program Scholarship (RTP). Figures were created with BioRender.com and ChemDraw.

## Table of Contents

This study introduces the covalent integration of enzymes in traditional gelatin bioinks for use as catalytic active scaffolding material for extrusion-based 3D bioprinting. Using horseradish peroxidase as a model enzyme, we achieve efficient post-printing photo-crosslinking and protein immobilization, while ensuring high enzyme stability, retention and prolonged catalytic activity. This novel approach enhances the bioactive properties in 3D-printed constructs and presents a crucial advancement for functional biofabricated tissues.

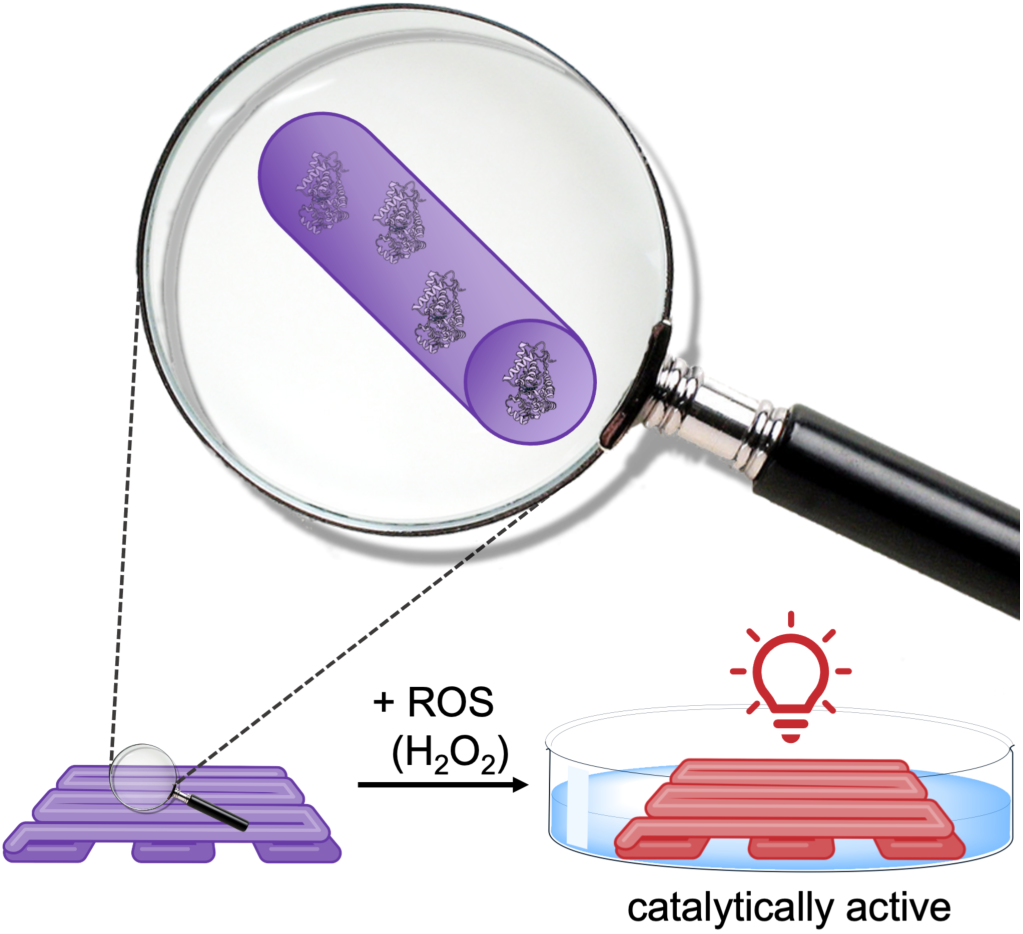

## References

1. Varkey, M.; Visscher, D. O.; van Zuijlen, P. P. M.; Atala, A.; Yoo, J. J. Skin bioprinting: the future of burn wound reconstruction? Burns & Trauma 2019, 7, 4.

2. Bian, L. Functional hydrogel bioink, a key challenge of 3D cellular bioprinting. APL Bioeng. 2020, 4, (3), 030401.

3. Mao, H. L.; Yang, L.; Zhu, H. F.; Wu, L. H.; Ji, P. H.; Yang, J. Q.; Gu, Z. W. Recent advances and challenges in materials for 3D bioprinting. Progress in Natural Science-Materials International 2020, 30, (5), 618–634.

4. Harley, W. S.; Li, C. C.; Toombs, J.; O’Connell, C. D.; Taylor, H. K.; Heath, D. E.; Collins, D. J. Advances in biofabrication techniques towards functional bioprinted heterogeneous engineered tissues: A comprehensive review. Bioprinting 2021, 23, e00147.

5. Bedell, M. L.; Navara, A. M.; Du, Y.; Zhang, S.; Mikos, A. G. Polymeric Systems for Bioprinting. Chem. Rev. 2020, 120, (19), 10744–10792.

6. Hoffman, A. S. Hydrogels for biomedical applications. Adv. Drug Delivery Rev. 2012, 64, 18–23.

7. Cui, X.; Li, J.; Hartanto, Y.; Durham, M.; Tang, J.; Zhang, H.; Hooper, G.; Lim, K.; Woodfield, T. Advances in Extrusion 3D Bioprinting: A Focus on Multicomponent Hydrogel-Based Bioinks. Adv Healthc Mater 2020, 9, (15), e1901648.

8. Unagolla, J. M.; Jayasuriya, A. C. Hydrogel-based 3D bioprinting: A comprehensive review on cell-laden hydrogels, bioink formulations, and future perspectives. Applied Materials Today 2020, 18, 100479.

9. Piao, Y.; You, H.; Xu, T.; Bei, H.-P.; Piwko, I. Z.; Kwan, Y. Y.; Zhao, X. Biomedical applications of gelatin methacryloyl hydrogels. Engineered Regeneration 2021, 2, 47–56.

10. Rajabi, N.; Rezaei, A.; Kharaziha, M.; Bakhsheshi-Rad, H. R.; Luo, H.; RamaKrishna, S.; Berto, F. Recent Advances on Bioprinted Gelatin Methacrylate-Based Hydrogels for Tissue Repair. Tissue Eng., Part A 2021, 27, (11-12), 679–702.

11. Liu, W.; Heinrich, M. A.; Zhou, Y.; Akpek, A.; Hu, N.; Liu, X.; Guan, X.; Zhong, Z.; Jin, X.; Khademhosseini, A.; Zhang, Y. S. Extrusion Bioprinting of Shear-Thinning Gelatin Methacryloyl Bioinks. Adv. Healthcare Mater. 2017, 6, (12), 1601451.

12. Kilic Bektas, C.; Hasirci, V. Cell Loaded GelMA:HEMA IPN hydrogels for corneal stroma engineering. J. Mater. Sci. Mater. Med. 2019, 31, (1), 2.

13. Montazerian, H.; Baidya, A.; Haghniaz, R.; Davoodi, E.; Ahadian, S.; Annabi, N.; Khademhosseini, A.; Weiss, P. S. Stretchable and Bioadhesive Gelatin Methacryloyl-Based Hydrogels Enabled by in Situ Dopamine Polymerization. ACS Appl. Mater. Interfaces 2021, 13, (34), 40290–40301.

14. Yang, Q.; Gao, B.; Xu, F. Recent Advances in 4D Bioprinting. Biotechnol. J. 2020, 15, (1), e1900086.

15. Leberfinger, A. N.; Dinda, S.; Wu, Y.; Koduru, S. V.; Ozbolat, V.; Ravnic, D. J.; Ozbolat, I. T. Bioprinting functional tissues. Acta Biomater. 2019, 95, 32–49.

16. Wang, R.; Wang, Y.; Yao, B.; Hu, T.; Li, Z.; Huang, S.; Fu, X. Beyond 2D: 3D bioprinting for skin regeneration. Int. Wound J. 2019, 16, (1), 134–138.

17. Howard, D.; Buttery, L. D.; Shakesheff, K. M.; Roberts, S. J. Tissue engineering: strategies, stem cells and scaffolds. J. Anat. 2008, 213, (1), 66–72.

18. Guebitz, G. M.; Nyanhongo, G. S. Enzymes as Green Catalysts and Interactive Biomolecules in Wound Dressing Hydrogels. Trends Biotechnol. 2018, 36, (10), 1040–1053.

19. Mandla, S.; Davenport Huyer, L.; Radisic, M. Review: Multimodal bioactive material approaches for wound healing. APL Bioeng. 2018, 2, (2), 021503.

20. Lee, Y.; Son, J. Y.; Kang, J. I.; Park, K. M.; Park, K. D. Hydrogen Peroxide-Releasing Hydrogels for Enhanced Endothelial Cell Activities and Neovascularization. ACS Appl. Mater. Interfaces 2018, 10, (21), 18372–18379.

21. Yang, K.; Han, Q.; Chen, B.; Zheng, Y.; Zhang, K.; Li, Q.; Wang, J. Antimicrobial hydrogels: promising materials for medical application. Int J Nanomedicine 2018, 13, 2217–2263.

22. Pinelli, F.; Magagnin, L.; Rossi, F. Progress in hydrogels for sensing applications: a review. Materials Today Chemistry 2020, 17, 100317.

23. Mandon, C. A.; Blum, L. J.; Marquette, C. A. Adding Biomolecular Recognition Capability to 3D Printed Objects. Anal. Chem. 2016, 88, (21), 10767–10772.

24. Wang, X.; Wang, Q. Enzyme-Laden Bioactive Hydrogel for Biocatalytic Monitoring and Regulation. Acc. Chem. Res. 2021, 54, (5), 1274–1287.

25. Labus, K. Effective detection of biocatalysts with specified activity by using a hydrogel-based colourimetric assay - beta-galactosidase case study. PLoS One 2018, 13, (10), e0205532.

26. Pose-Boirazian, T.; Martinez-Costas, J.; Eibes, G. 3D Printing: An Emerging Technology for Biocatalyst Immobilization. Macromol. Biosci. 2022, 22, (9), e2200110.

27. Remonatto, D.; Izidoro, B. F.; Mazziero, V. T.; Catarino, B. P.; do Nascimento, J. F. C.; Cerri, M. O.; Andrade, G. S. S.; Paula, A. V. d. 3D printing and enzyme immobilization: An overview of current trends. Bioprinting 2023, 33, e00289.

28. Schmieg, B.; Dobber, J.; Kirschhofer, F.; Pohl, M.; Franzreb, M. Advantages of Hydrogel-Based 3D-Printed Enzyme Reactors and Their Limitations for Biocatalysis. Front Bioeng Biotechnol 2018, 6, 211.

29. Schneider, K. P.; Gewessler, U.; Flock, T.; Heinzle, A.; Schenk, V.; Kaufmann, F.; Sigl, E.; Guebitz, G. M. Signal enhancement in polysaccharide based sensors for infections by incorporation of chemically modified laccase. N. Biotechnol. 2012, 29, (4), 502–509.

30. Baretta, R.; Gabrielli, V.; Frasconi, M. Nanozyme-Cellulose Hydrogel Composites Enabling Cascade Catalysis for the Colorimetric Detection of Glucose. Acs Applied Nano Materials 2022, 5, (10), 13845–13853.

31. Abdel-Mageed, H. M.; Abd El Aziz, A. E.; Abdel Raouf, B. M.; Mohamed, S. A.; Nada, D. Antioxidant-biocompatible and stable catalase-based gelatin-alginate hydrogel scaffold with thermal wound healing capability: immobilization and delivery approach. 3 Biotech 2022, 12, (3), 73.

32. Kurahashi, T.; Fujii, J. Roles of Antioxidative Enzymes in Wound Healing. J Dev Biol 2015, 3, (2), 57–70.

33. Molley, T. G.; Jiang, S.; Ong, L.; Kopecky, C.; Ranaweera, C. D.; Jalandhra, G. K.; Milton, L.; Kardia, E.; Zhou, Z.; Rnjak-Kovacina, J.; Waters, S. A.; Toh, Y. C.; Kilian, K. A. Gas-modulating microcapsules for spatiotemporal control of hypoxia. Proc. Natl. Acad. Sci. U. S. A. 2023, 120, (16), e2217557120.

34. Yeroslavsky, G.; Girshevitz, O.; Foster-Frey, J.; Donovan, D. M.; Rahimipour, S. Antibacterial and antibiofilm surfaces through polydopamine-assisted immobilization of lysostaphin as an antibacterial enzyme. Langmuir 2015, 31, (3), 1064–1073.

35. Kim, S.; Fan, J.; Lee, C. S.; Lee, M. Dual Functional Lysozyme-Chitosan Conjugate for Tunable Degradation and Antibacterial Activity. ACS Appl Bio Mater 2020, 3, (4), 2334–2343.

36. Wenger, L.; Radtke, C. P.; Gopper, J.; Worner, M.; Hubbuch, J. 3D-Printable and Enzymatically Active Composite Materials Based on Hydrogel-Filled High Internal Phase Emulsions. Front. Bioeng. Biotechnol. 2020, 8, 713.

37. Shen, J.; Zhang, S.; Fang, X.; Salmon, S. Advances in 3D Gel Printing for Enzyme Immobilization. Gels 2022, 8, (8), 460.

38. Meyer, J.; Meyer, L. E.; Kara, S. Enzyme immobilization in hydrogels: A perfect liaison for efficient and sustainable biocatalysis. Eng. Life Sci. 2022, 22, (3-4), 165–177.

39. Ferracci, G.; Zhu, M.; Ibrahim, M. S.; Ma, G.; Fan, T. F.; Lee, B. H.; Cho, N. J. Photocurable Albumin Methacryloyl Hydrogels as a Versatile Platform for Tissue Engineering. ACS Appl Bio Mater 2020, 3, (2), 920–934.

40. Yokoyama, K.; Nio, N.; Kikuchi, Y. Properties and applications of microbial transglutaminase. Appl. Microbiol. Biotechnol. 2004, 64, (4), 447–454.

41. Sakai, S.; Mochizuki, K.; Qu, Y.; Mail, M.; Nakahata, M.; Taya, M. Peroxidase-catalyzed microextrusion bioprinting of cell-laden hydrogel constructs in vaporized ppm-level hydrogen peroxide. Biofabrication 2018, 10, (4), 045007.

42. Choi, S.; Ahn, H.; Kim, S. H. Tyrosinase-mediated hydrogel crosslinking for tissue engineering. J. Appl. Polym. Sci. 2021, 139, (14), 51887.

43. Schmieg, B.; Schimek, A.; Franzreb, M. Development and performance of a 3D-printable poly(ethylene glycol) diacrylate hydrogel suitable for enzyme entrapment and long-term biocatalytic applications. Eng. Life Sci. 2018, 18, (9), 659–667.

44. Shao, Y.; Liao, Z.; Gao, B.; He, B. Emerging 3D Printing Strategies for Enzyme Immobilization: Materials, Methods, and Applications. ACS Omega 2022, 7, (14), 11530–11543.

45. Wenger, L.; Radtke, C. P.; Gerisch, E.; Kollmann, M.; Niemeyer, C. M.; Rabe, K. S.; Hubbuch, J. Systematic evaluation of agarose- and agar-based bioinks for extrusion-based bioprinting of enzymatically active hydrogels. Front Bioeng Biotechnol 2022, 10, 928878.

46. Gkantzou, E.; Weinhart, M.; Kara, S. 3D printing for flow biocatalysis. RSC Sustainability 2023, 1, (7), 1672–1685.

47. Pinyakit, Y.; Romphophak, P.; Painmanakul, P.; Hoven, V. P. Introduction of an Ambient 3D-Printable Hydrogel Ink to Fabricate an Enzyme-Immobilized Platform with Tunable Geometry for Heterogeneous Biocatalysis. Biomacromolecules 2023, 24, (7), 3138–3148.

48. Devillard, C. D.; Mandon, C. A.; Lambert, S. A.; Blum, L. J.; Marquette, C. A. Bioinspired Multi-Activities 4D Printing Objects: A New Approach Toward Complex Tissue Engineering. Biotechnol. J. 2018, 13, (12), e1800098.

49. Smith, P. T.; Narupai, B.; Tsui, J. H.; Millik, S. C.; Shafranek, R. T.; Kim, D. H.; Nelson, A. Additive Manufacturing of Bovine Serum Albumin-Based Hydrogels and Bioplastics. Biomacromolecules 2020, 21, (2), 484–492.

50. Ying, G. L.; Jiang, N.; Maharjan, S.; Yin, Y. X.; Chai, R. R.; Cao, X.; Yang, J. Z.; Miri, A. K.; Hassan, S.; Zhang, Y. S. Aqueous Two-Phase Emulsion Bioink-Enabled 3D Bioprinting of Porous Hydrogels. Adv. Mater. 2018, 30, (50), e1805460.

51. Asim, S.; Tabish, T. A.; Liaqat, U.; Ozbolat, I. T.; Rizwan, M. Advances in Gelatin Bioinks to Optimize Bioprinted Cell Functions. Adv Healthc Mater 2023, 12, (17), e2203148.

52. Shirahama, H.; Lee, B. H.; Tan, L. P.; Cho, N. J. Precise Tuning of Facile One-Pot Gelatin Methacryloyl (GelMA) Synthesis. Sci. Rep. 2016, 6, (1), 31036.

53. Lee, B. H.; Shirahama, H.; Cho, N. J.; Tan, L. P. Efficient and controllable synthesis of highly substituted gelatin methacrylamide for mechanically stiff hydrogels. RSC Adv. 2015, 5, (128), 106094–106097.

54. Konhauser, M.; Kannaujiya, V. K.; Kaltbeitzel, J.; Winterwerber, P.; Bohm, E.; Breitenbach, B.; Wich, P. R. Dual-Responsive Enzyme-Polysaccharide Conjugate as a Nanocarrier System for Enzyme Prodrug Therapy. Biomacromolecules 2023, 24, (5), 2138–2148.

55. Mishin, V.; Gray, J. P.; Heck, D. E.; Laskin, D. L.; Laskin, J. D. Application of the Amplex red/horseradish peroxidase assay to measure hydrogen peroxide generation by recombinant microsomal enzymes. Free Radical Biol. Med. 2010, 48, (11), 1485–1491.

56. Zhang, Y.; Schmid, Y. R. F.; Luginbuhl, S.; Wang, Q.; Dittrich, P. S.; Walde, P. Spectrophotometric Quantification of Peroxidase with p-Phenylene-diamine for Analyzing Peroxidase-Encapsulating Lipid Vesicles. Anal. Chem. 2017, 89, (10), 5484–5493.

57. Azevedo, A. M.; Martins, V. C.; Prazeres, D. M.; Vojinovic, V.; Cabral, J. M.; Fonseca, L. P. Horseradish peroxidase: a valuable tool in biotechnology. Biotechnol. Annu. Rev. 2003, 9, 199–247.

58. Bortolami, S.; Cavallini, L. Enhanced detection of H2O2 in cells expressing horseradish peroxidase. Free Radical Res. 2009, 43, (5), 446–456.

59. Navapour, L.; Mogharrab, N.; Amininasab, M. How modification of accessible lysines to phenylalanine modulates the structural and functional properties of horseradish peroxidase: a simulation study. PLoS One 2014, 9, (10), e109062.

60. Fairbanks, B. D.; Schwartz, M. P.; Bowman, C. N.; Anseth, K. S. Photoinitiated polymerization of PEG-diacrylate with lithium phenyl-2,4,6-trimethylbenzoylphosphinate: polymerization rate and cytocompatibility. Biomaterials 2009, 30, (35), 6702–6707.

61. Branco, M. C.; Pochan, D. J.; Wagner, N. J.; Schneider, J. P. The effect of protein structure on their controlled release from an injectable peptide hydrogel. Biomaterials 2010, 31, (36), 9527–9534.

62. Zhou, M.; Diwu, Z.; Panchuk-Voloshina, N.; Haugland, R. P. A stable nonfluorescent derivative of resorufin for the fluorometric determination of trace hydrogen peroxide: applications in detecting the activity of phagocyte NADPH oxidase and other oxidases. Anal. Biochem. 1997, 253, (2), 162–168.

63. Kim, N. H.; Jeong, M. S.; Choi, S. Y.; Kang, J. H. Peroxidase activity of cytochrome. Bull. Korean Chem. Soc. 2004, 25, (12), 1889–1892.

