## Supporting Information for "Enzyme Bioink for the 3D Printing of Biocatalytic Materials"

### Materials

Gelatin (300 bloom, Type A), methacrylic anhydride (MAA), fluorescamine, horseradish peroxidase (Type II, 150-200 U/mg), deuterium oxide ($D_2O$), 2,2'-Azino-bis(3-ethylbenzothiazoline-6-sulfonic acid) diammonium salt (ABTS) and Dulbecco′s Modified Eagle′s Medium - high glucose (DMEM) were purchased from Sigma-Aldrich; collagenase (type I) was purchased from Thermo Fisher Australia; lithium phenyl-2,4,6-trimethylbenzoylphosphinate (LAP) from Ambeed, hydrogen peroxide (27-30%), dimethyl sulfoxide (DMSO) from Chem-Supply Pty Ltd.

### Analytical Instrumentation

$^1$H-NMR spectra were recorded using a 400 MHz Bruker Topspin Fourier spectrometer. Infrared (IR) spectra were recorded using a Nicolet Avatar 330-IR (ATR-unit) spectrometer. Anton Paar MCR 302 rheometer with a parallel plate geometry (25 mm disk, 1 mm measuring distance, 600 $\mu$L of liquid sample) was used for rheological tests. Confocal microscopy images were captured using a Zeiss LSM 900 CD7 and Zeiss LSM 800 microscope

### Gel-MA Characterization ($^1$H-NMR)

To determine the changes in gelatin structure due to methacrylation, proton nuclear magnetic resonance ($^1$H-NMR) was performed. A 0.5 mL sample of 50 mg/mL Gel-MA and gelatin samples were prepared by dissolving 25 mg of each in 0.5 mL of $D_2O$ at 40 °C in an incushaker. The samples were then transferred to clean NMR glass vials and analyzed in a 400 MHz Bruker Topspin Fourier spectrometer. The designated peaks in the spectra indicate the change in the molecular structure due to methacrylate groups.

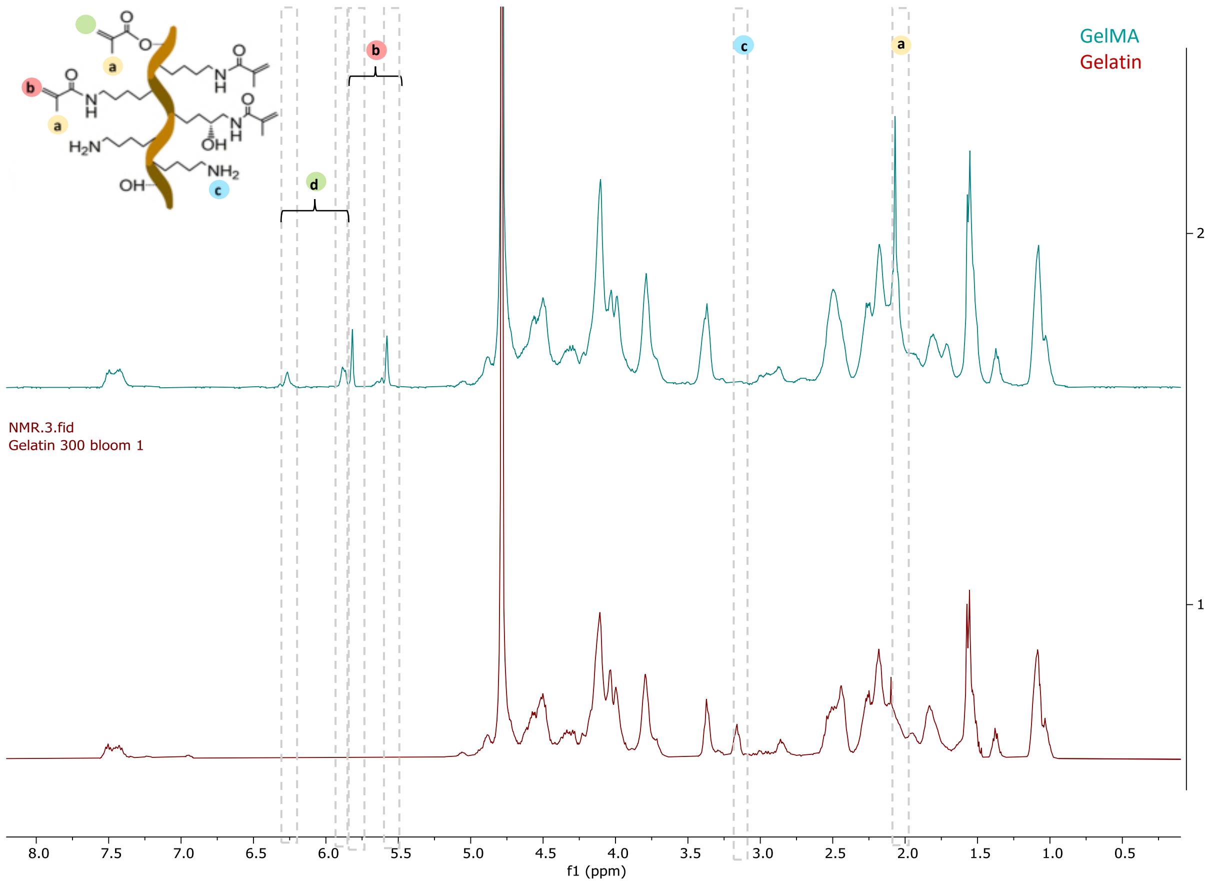

*Figure S1. $^1$H-NMR spectra in $D_2O$ at 37 °C (400 MHz) for comparison of (red) Gelatin and Gel-MA (blue). Spectrum a shows the characteristic signal of the  protons (H-g, H-h)*

**Gel-MA Characterization (Fluorescamine Assay)**

Gelatin and Gel-MA were adjusted to a concentration of 1.6 mg/mL via absorbance measurement at 280 nm. Gelatin was used as an external standard in a concentration range of 0.016 mg/mL to 2 mg/mL. For the assay, 125 $\mu$L PBS (0.01 M, pH 7.4), as well as 25 $\mu$L sample/standard, was added in triplicates into each well of a 96-black-well-microplate (flat bottom). Just before the measurement, 50 $\mu$L of 0.3 mg/mL freshly prepared fluorescamine solution in DMSO was mixed in before being analyzed with a fluorescence plate reader (Ex/Em = 380/460 nm); The degree of substitution (DS) of amines to methacrylate groups was calculated as follows:

$$\text{DS} = \left(1 - \frac{\text{amine content in Gel-MA or HRP-MA}\left[\frac{\text{mmol}}{\text{g}}\right]}{\text{amine content in gelatin or HRP}\left[\frac{\text{mmol}}{\text{g}}\right]}\right) \cdot 100\% \qquad \textit{Eq 1}$$

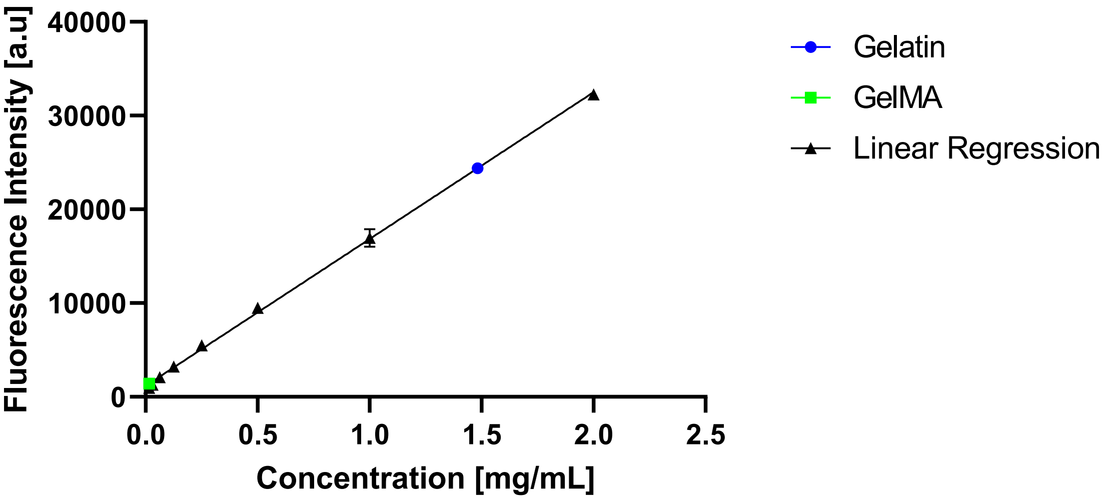

*Figure S2. Standard curve to determine the degree of substitution of Gel-MA*

**HRP-MA Characterization (FTIR Spectroscopy)**

As a verification method for successful methacrylation of HRP-MA, Fourier-transform infrared spectroscopy was carried out. Here the FTIR spectrum of HRP-MA was compared with the FTIR spectrum of HRP. Additional peaks at wavenumber bands at 3112 $cm^{-1}$ (CH stretching at the $CH_2$ group), at 1046 $cm^{-1}$ (C–O–C stretching) and a strong peak at 1638 $cm^{-1}$ (C=O) confirm methacrylation on the enzyme molecule structure.

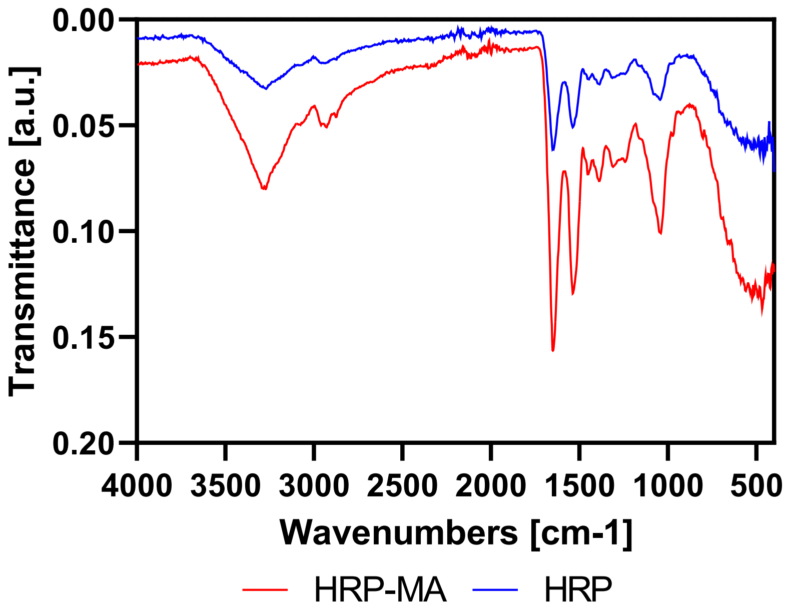

*Figure S3. FTIR spectra of HRP (blue) and HRP-MA (red)*

**Rheological Measurements - Shear Sweep Tests**

Shear sweep tests were performed on all the hydrogels with a log ramp-up rate from 0.02% shear strain up to 200% at 1 Hz frequency over 8 minutes to determine the yield stress.

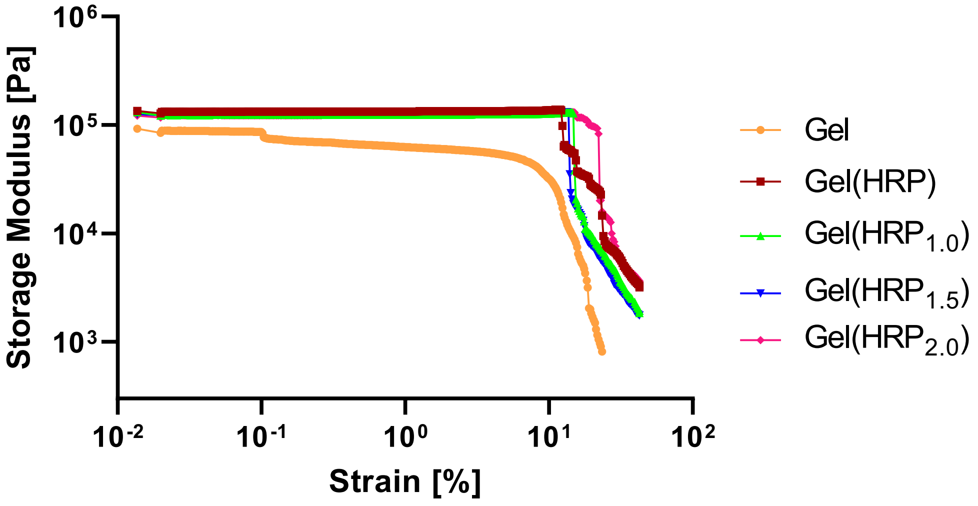

*Figure S4. Shear strain amplitude sweeps of different hydrogels*

**Rheological Measurements - Storage Modulus as a Function of Time with UV Light**

We were able to tune the storage modulus (stiffness) of our Gel(HRP) hydrogel by adjusting the UV exposure time from 5 seconds to 15 seconds. The experimental setup and conditions were identical to those described in the "Rheology of Bioink" section of the Materials and Methods, with the exception that the test duration was reduced to 10 minutes instead of 20 minutes, and the UV light exposure was initiated at the 3rd minute instead of the 4th minute.

Upon analysis, we observed that the hydrogel exhibited storage modulus values of approximately 30 kPa, 55 kPa, and 85 kPa when the UV light exposure time was set to 5, 10, and 15 s, respectively. This finding demonstrates our capability to finely control and tune the stiffness of the hydrogels, which holds significant promise for various applications.

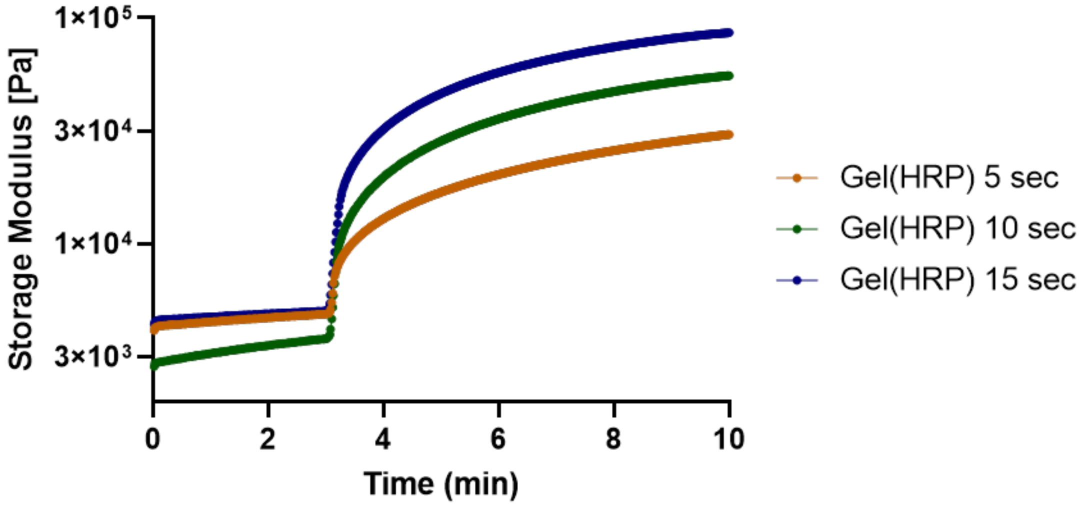

*Figure S5. Effect of varying UV light exposure time on the storage modulus (G') of Gel(HRP) hydrogel.*

**Enzyme Washing Test and Verification of Equal Initial Enzyme Amounts**

The enzyme bioink containing either the native HRP (Gel(HRP$_{native}$)) or the methacrylated HRP (Gel(HRP)) was pipetted in 20 $\mu$L droplets at 37 °C into separate wells of a 12-well cell culture plate (CoStar, 3513) respectively in triplicates. Subsequently, these droplets were photo-crosslinked at 395 nm for 1, 3 or 5 min to stable gel droplets, and immersed with 2 mL DMEM media. As a control, 20 $\mu$L of enzyme bioink was pipetted directly into a 2 mL media, mixed for full dissolution, and then treated equally with UV light (395 nm), shaking at 80 rpm and 37 °C and 5 min of sonication as the crosslinked stable droplets. Then, 10 $\mu$L of the media was transferred into a 96-well plate (Sarstaedt) to determine the containing enzyme activity via ABTS assay. Therefore, 40 $\mu$L ddH$_2$O, 100 $\mu$L ABTS solution, and 50 $\mu$L of $H_2O_2$ (stock solution 0.1 mM in ddH$_2$O) were added additionally and the absorbance was measured immediately at 410 nm for 3 min every 10 s in a kinetic cycle. The analysis was performed via GraphPad Prism 9 where the initial rate of the absorbance increase of each droplet was compared and the controls normalized to 100%.

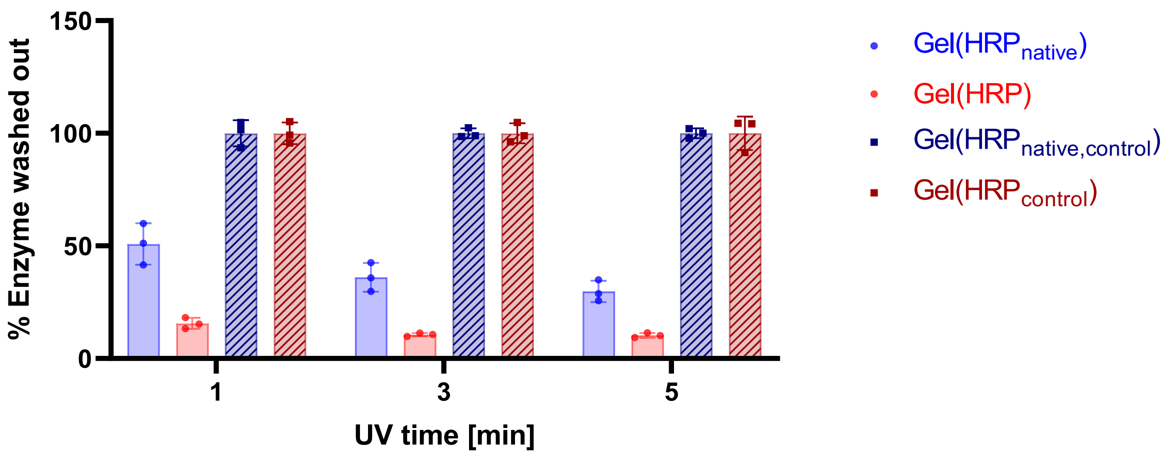

*Figure S6. Effect of varying UV light exposure time on the enzyme retention of Gel(HRP) and Gel(HRP$_{native}$). Control droplets with non-crosslinked samples confirm that all initial droplets contain the same total enzyme amount/activity. This value was set to 100% enzyme activity.*

**Collagenase Degradation Assay**

The two enzyme bioinks containing either the native HRP (Gel($HRP_{native}$)) or the methacrylated HRP (Gel(HRP)) were pipetted in 20 $\mu$L droplets at 37 °C into separate wells of a 12-well cell culture plate (CoStar, 3513) respectively in triplicates. Subsequently, these droplets were photo-crosslinked at 395 nm for 1 min, and immersed with 2 mL collagenase solution (0.35 mg/mL). As control for the starting enzyme activity, the same amount of 20 $\mu$L of enzyme bioink was pipetted directly into a 2 mL of collagenase solution, mixed for full dissolution, and then treated equally. After full degradation of the droplets after 3 h of incubation at 37 °C, the enzyme activity of the solution was measured. Therefore, 10 $\mu$L of the sample solution, 40 $\mu$L dd$H_2O$, 100 $\mu$L ABTS solution, and 50 $\mu$L of $H_2O_2$ (stock solution 0.1 mM in dd$H_2O$) has been added into a into a 96-well plate (Sarstaedt). Subsequently, the absorbance was measured at 410 nm for 3 min every 10 s in a kinetic cycle. The analysis was performed via GraphPad Prism 9 where the initial rate of the absorbance increase of each triplicate was compared and the controls normalized to 100%.

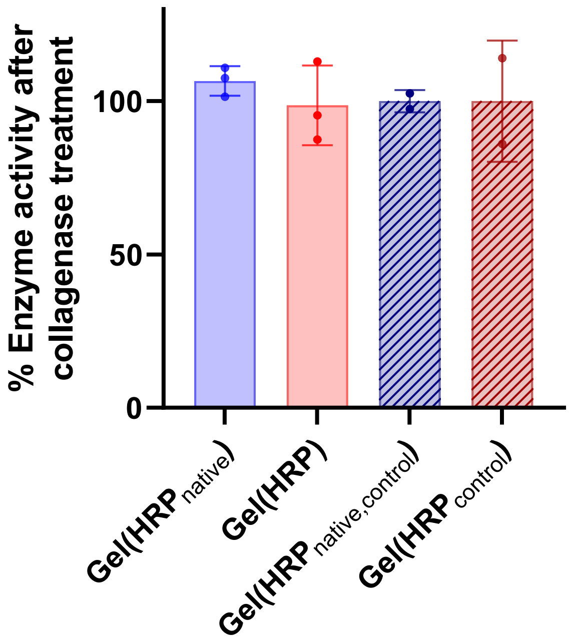

*Figure S7. Collagenase degradation assay to determine the enzyme activity after droplet formation, photocrosslinking, and collagenase degradation of hydrogel.*

**ABTS Assay on Cell-laden Hydrogels**

Gel($HRP_{native}$) and Gel(HRP) hydrogels underwent ABTS testing following incubation at various time intervals with cells. These hydrogels were immersed in solutions containing ABTS and $H_2O_2$. In the presence of HRP within the hydrogels, a reaction occurred creating a purple azo color formation in hydrogels with HRP within. The results, as illustrated in the figure, unequivocally demonstrate that methacrylated HRP remains detectable within the hydrogels even after 7 days of co-incubation with cells. In stark contrast, $HRP_{native}$ exhibited minimal presence within the hydrogels. These findings affirm not only the successful crosslinking of methacrylated HRP with the hydrogel but also its remarkable stability, persisting for more than 7 days even in a cellular environment.

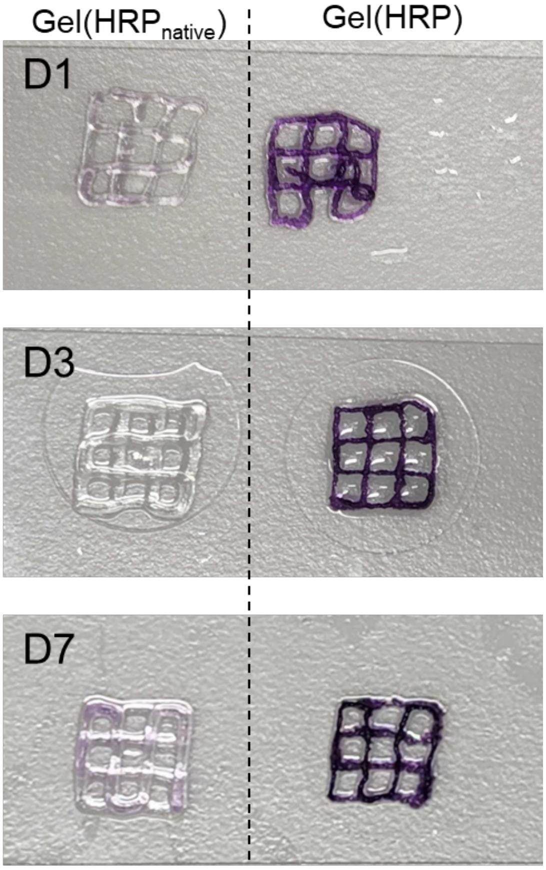

*Figure S8. Photographs of 3D printed grids of cell-laden Gel($HRP_{native}$) and Gel(HRP) over days 1, 3, and 7 after reaction with ABTS reagent.*

**Enzyme Bioink Activity Test Inside Gel (Additional Experiments)**

For all samples, a stock solution of 11 w/v % Gel-MA was prepared at 40 °C. Next, LAP (2.5 mg/mL) and enzyme (0.625 mg/mL for Gel($HRP_{native}$) and Gel(HRP), and 0.3125 mg/mL for Gel($HRP_{50\%}$)) were added in the absence of light. To avoid contamination, the hydrogel was filtered once with a syringe filter (33 mm, Ø 0.45 µm) before further use. Droplets of 10 $\mu$l were pipetted at 37 °C into separate wells of a 12-well cell culture plate (CoStar, 3513) and subsequently photo-crosslinked at 395 nm (40 mW $cm^{-2}$) for 1 min to stable gel droplets. Then these droplets were immersed with 2 mL DMEM media and incubated while shaking at 80 rpm at 37 ºC with additional sonication for 5 min. After 24 hours of washing in media, the activity inside the gels was visualized via confocal microscopy through the oxidative conversion of Amplex Red to resorufin after the addition of $H_2O_2$. Tile images of each enzyme bioink droplet (Gel($HRP_{native}$), Gel(HRP), Gel($HRP_{50\%}$)) of identical size and z-height from the plate bottom were visualized (objective 5x, NA 0.35; Ex/Em = 568/581 nm) by comprising 8 slices of z-stack covering the full droplet.

Before imaging, the media was removed, and the droplets were rinsed once with PBS. 1 mL of PBS (0.01 M, pH 7.4) was added to each of the wells containing the bioink droplets. Then, 10 $\mu$L of Amplex Red solution (stock solution 10 mM in DMSO) was added, mixed, and incubated for 5 min before adding 5 $\mu$L of $H_2O_2$ (stock solution 0.1 mM in dd$H_2O$) with an incubation time of 1 min. The solution was discarded, and any excess was removed gently with lint-free wipes. A 10 min timeframe was allowed for penetration before imaging. Image analysis was performed with Imaris software (version 9.9.0). The fluorescence intensity was calculated with ImageJ by selecting the droplet area of each plane and carrying out a mean measurement which was then summed up to a total value.

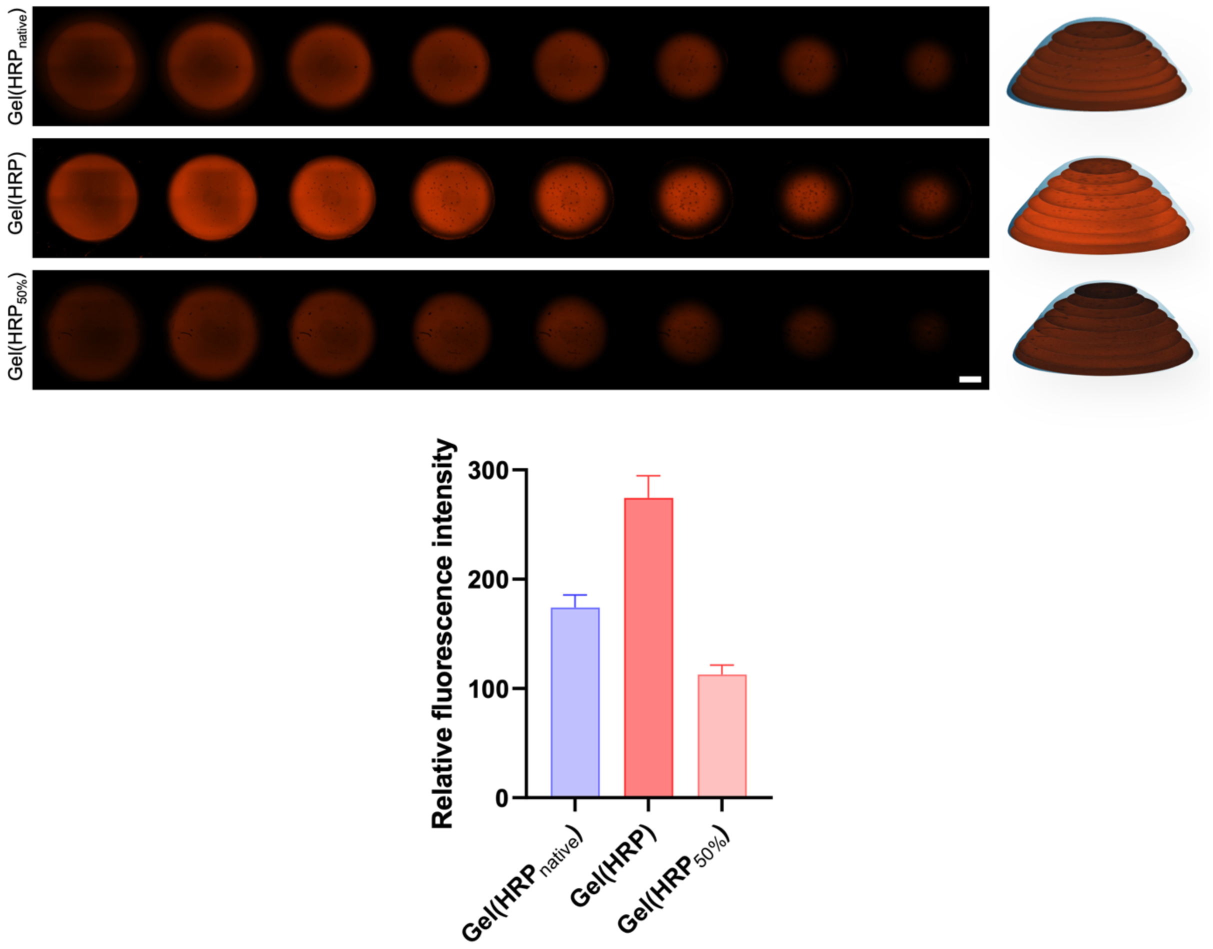

*Figure S9. Enzyme bioink activity test inside 10 µL hydrogel droplets. Confocal images of z-stack plane images of different enzyme bioink droplets Gel($HRP_{native}$), Gel(HRP), and Gel($HRP_{50\%}$). After washing for 24 h the droplets were treated with Amplex Red and $H_2O_2$ and images were recorded with a Zeiss CD7 LSM 900 (Ex/Em = 568/581 nm); Scale bar = 1 mm; Right: z-stack plane images reassembled into 3D droplets. Bottom: ImageJ analysis of mean fluorescence intensity in each layer of the respective droplet; Error bars represent standard deviation obtained by ImageJ.*

**Cell Culture - Average Cell Area**

To determine the average cell area in the hydrogels, we utilized Fiji Image J software, measuring the individual cell size and coverage in each image. Five images were captured for each hydrogel sample, and the area covered by each cell in these images was recorded and subsequently averaged. The objective of the experiment is to assess whether the cells exhibit an increase in their size and merge, resulting in a greater occupied area over time.

Overall, the analysis of the hydrogels from day 1 (D1) to day 7 (D7) revealed a slight increase in the average cell area. Figure S10(A) compares the hydrogels with and without our enzyme, HRP-MA, while Figure S10(B) shows the outcome with a progressive increase in our enzyme concentration within these hydrogels. This observation indicates improved cell viability and growth within the hydrogels.

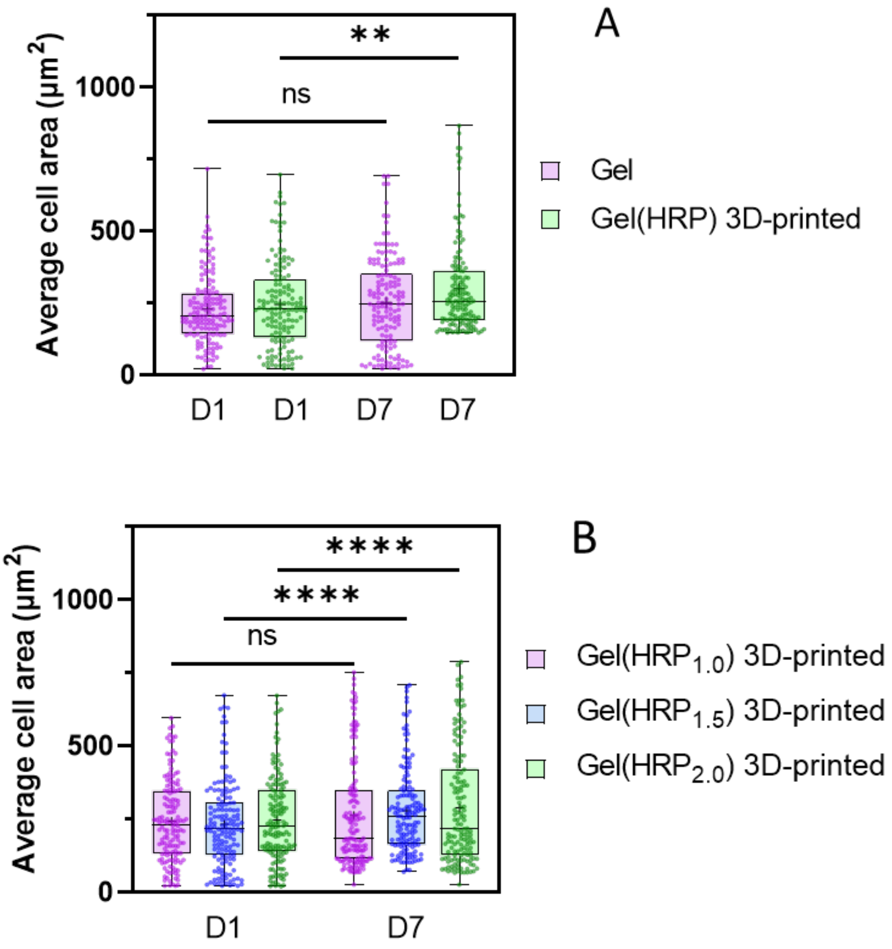

*Figure S10. (A) Average cell area of ADSCs in Gel and Gel(HRP) hydrogels. (B) The average cell area of ADSCs in Gel($HRP_{1.0}$) 3D-printed, Gel($HRP_{1.5}$) 3D-printed, and Gel($HRP_{2.0}$) 3D-printed hydrogels; the mean of the average cell area is represented by a small plus sign (+); n=150, number of cells; **$P < 0.01$, ****$P < 0.0001$.*
